# Polyglutamine-expanded androgen receptor disrupts muscle triad, calcium dynamics and the excitation-contraction coupling gene expression program

**DOI:** 10.1101/618405

**Authors:** M Chivet, C Marchioretti, M Pirazzini, D Piol, C Scaramuzzino, JM Polanco, S Nath, E Zuccaro, L Nogara, M Canato, L Marcucci, S Parodi, V Romanello, A Armani, M D’Antonio, F Sambataro, E Dassi, E Pegoraro, G Sorarù, C Rinaldi, AP Lieberman, B Blaauw, M Sandri, M Basso, M Pennuto

## Abstract

Spinal and bulbar muscular atrophy (SBMA) is caused by polyglutamine (polyQ) expansions in the androgen receptor (*AR*) gene. Although clinical and experimental evidence highlight a primary role for skeletal muscle in the onset, progression, and outcome of disease, the pathophysiological and molecular processes underlying SBMA muscle atrophy are poorly understood. Here we show that polyQ-expanded AR alters intrinsic muscle force generation before denervation. Reduced muscle force was associated with a switch in fiber-type composition, disrupted muscle striation, altered calcium (Ca^++^) dynamics in response to muscle contraction, and aberrant expression of excitation-contraction coupling (ECC) machinery genes in transgenic, knock-in and inducible SBMA mice and patients. Importantly, treatment to suppress polyQ-expanded AR toxicity restored ECC gene expression back to normal. Suppression of AR activation by surgical castration elicited similar ECC gene expression changes in normal mice, suggesting that AR regulates the expression of these genes in physiological conditions. Bioinformatic analysis revealed the presence of androgen-responsive elements on several genes involved in muscle function and homeostasis, and experimental evidence showed AR-dependent regulation of expression and promoter occupancy of the most up-regulated gene from transcriptomic analysis in SBMA muscle, i.e. sarcolipin, a key ECC gene. These observations reveal an unpredicted role for AR in the regulation of expression of genes involved in muscle contraction and Ca^++^ dynamics, a level of muscle function regulation that is disrupted in SBMA muscle, yet restored by pharmacologic treatment.

## Introduction

Expansions of the polyQ-encoding CAG trinucleotide tandem repeat over 38 in exon 1 of the *AR* gene leads to SBMA, also known as Kennedy’s disease (1). SBMA belongs to the family of polyQ diseases, which includes Huntington’s disease, dentatorubral-pallidoluysian atrophy, and six types of spinocerebellar ataxia (2, 3). SBMA is an X-linked adult-onset neuromuscular disease characterized by the progressive dysfunction and degeneration of lower motor neurons (MNs) and skeletal muscle weakness, fasciculations and atrophy (4). In addition, patients develop signs of androgen insensitivity syndrome, such as gynecomastia, erectile dysfunction and reduced fertility (5, 6). Moreover, patients may present signs of metabolic syndrome, such as insulin resistance, glucose intolerance, hyperlipidemia and non-alcoholic fatty liver (6–8). Among polyQ diseases, a unique feature of SBMA is sex-specificity with males primarily affected, and females showing only mild disease manifestations even if homozygous for the mutation (9). Males are affected because they have high levels of circulating testosterone in the serum (10). Neuromuscular phenotype, sex-dependency, metabolic alterations and peripheral symptoms are well recapitulated in transgenic and knock-in mouse models of disease (11–14). Mechanistically, the mild signs of androgen insensitivity and endocrine abnormalities implicate partial AR loss of function (LOF) underlying disease pathogenesis (5, 6). However, pure AR LOF mutations lead to different degrees of androgen insensitivity syndrome in the absence of neurological symptoms, indicating that SBMA is not a pure LOF disease. Rather, SBMA results from partial AR LOF in combination with polyQ-AR neomorphic gain of function (GOF) (15). A key factor contributing to the toxic GOF of polyQ-AR is the deposition of the disease protein into micro-aggregates and inclusion bodies, and specific types of polyQ-AR-positive aggregates have been linked to neurotoxicity *in vitro* and *in vivo* (12, 16–19). Although the androgen-dependent nature of disease and experimental evidence support chemical and physical castration as a therapeutic strategy for SBMA (10, 11, 13), clinical trials based on this approach showed benefits limited to a subset of patients (20–22).

In the last decade clinical and experimental evidence have established that in SBMA skeletal muscle atrophy is not only secondary to MN dysfunction, but it also results from primary cell-autonomous toxicity of polyQ-expanded AR in this peripheral tissue (23, 24). Early muscle pathology and clear signs of myopathy are present in patients, including elevated levels of serum creatine kinase (CK), myofiber degeneration and structures similar to central cores detected before clinical symptoms (25–28). Likewise, signs of myopathy and reduced intrinsic muscle force precede spinal cord pathology also in transgenic and knock-in mouse models of SBMA (11, 29). Interestingly, fast-twitch muscles were mainly affected and displayed a glycolytic-to-oxidative fiber-type switch with alterations in lipid homeostasis regardless of denervation (30). Overexpression of a non-expanded AR selectively in muscle elicited a phenotype that for several aspects resembles SBMA (31), whereas ablation of polyQ-expanded AR expression selectively in muscle prevented disease manifestations (32), indicating that deregulation of androgen signaling in muscle is necessary and sufficient to cause functional alterations of the motor unit. The relevance of skeletal muscle in SBMA is underlined by two clinically relevant aspects. First, blood biomarkers of muscle damage correlated with disease severity (33). Second, skeletal muscle has been shown to be a valuable target tissue for therapy development. Pharmacologic intervention to either silence polyQ-expanded AR expression with antisense oligonucleotides (ASOs) in peripheral tissues, specially skeletal muscle (34), or to promote its degradation and stimulate muscle hypertrophy with the insulin-like growth factor 1 and the beta-agonist clenbuterol (35, 36) ameliorated disease manifestations, including metabolic defects in muscle (14). These observations further support the idea that not only the MN, but also the skeletal muscle is central to disease and represents a key target-tissue for therapy development. However, the mechanism through which deregulation of androgen signaling leads to muscle weakness and atrophy is poorly understood. Here, we provide the first evidence that polyQ expansions in the AR disrupt the ECC machinery and alter Ca^++^ dynamics in skeletal muscle. Strikingly, these pathological processes occurred before denervation, preceded motor dysfunction, were detected upon both chronic and acute induction of mutant AR expression and were found in patients with SBMA. Pharmacologic treatment to suppress polyQ-expanded AR toxicity successfully restored ECC gene expression changes. Importantly, we provide evidence that AR regulates the expression of specific ECC genes, a level of regulation of gene expression that is lost in SBMA.

## Results

### AR100Q mice display age-dependent AR aggregation, inclusion body formation, body weight loss, motor dysfunction and premature death

Here we sought to elucidate the mechanism through which polyQ expansions in AR cause muscle atrophy. To model SBMA, we generated transgenic mice expressing human AR (hAR) with an expanded polyQ tract (AR100Q) under a promoter for ubiquitous and constitutive expression in mammalian systems (**Supplementary Figure 1**). We first verified that AR100Q mice express physiological levels of monomeric hAR transgene in tissues that primarily degenerate in SBMA, i.e. spinal cord, brainstem, and skeletal muscle (**Figure 1A**). By Western blotting and filter retardation assay we found accumulation of high-molecular weight (HMW) species and micro-aggregates in the brainstem and skeletal muscle of 8-week-old AR100Q mice (**Figure 1A and Supplementary Figure 2A-D**). AR aggregation and inclusion body formation have been extensively characterized in neurons, but not in muscle (12, 16–19). In skeletal muscle polyQ-expanded AR aggregation was 2% sodium dodecyl sulphate (SDS)-resistant and was detected as early as 4 weeks of age mainly in muscles composed of both fast-glycolytic and slow-oxidative fibers and that degenerate in SBMA, such as quadriceps, gastrocnemius, and tibialis anterior (TA), as well as muscles composed of fast-glycolytic fibers, such as extensor digitorum longus (EDL), and to a lower extent in muscles composed of slow-oxidative fibers, such as soleus, and diaphragm and heart, which are only mildly affected in SBMA, indicating that polyQ-expanded AR aggregation occurs early and is affected by muscle metabolism. No polyQ-expanded AR aggregation was detected in other peripheral tissues, such as testis and adrenal glands that express levels of AR similar to muscle. Moreover, the amount of aggregated polyQ-expanded AR with respect to monomeric AR was higher in male compared to female AR100Q mice, suggesting that aggregation is enhanced by androgens (**Supplementary Figure 2E**). By immuno-histochemical and confocal microscopy analysis in intact fibers isolated from different skeletal muscles of 8-week-old AR100Q mice, polyQ-expanded AR showed a distinct punctate nuclear pattern (**Figure 1B and Supplementary Figures 3 and 4A-D**). Formation of muscle intranuclear inclusions changed with age. PolyQ-expanded AR was diffused in the myonuclei of 15-day-old AR100Q mice. Notably, we observed several small AR-positive puncta in the myonuclei at 4 weeks of age, when serum testosterone levels physiologically increase in mice, and less yet larger inclusion bodies per nucleus at 8 weeks of age. Nuclear inclusions were detected also in female mice, yet to a much lower extent compared to male mice, suggesting that the process of AR-positive inclusion formation is enhanced by androgens. No inclusions were detected in smooth muscles, such as bladder (**Supplementary Figure 4E**). Rather, inclusions were detected in several striated muscles, to a higher extent in EDL and flexor digitorum brevis (FDB) than soleus (**Supplementary Figure 4F**). Inclusion bodies were detected both in sub-synaptic and non-sub-synaptic nuclei (**Supplemental Figure 5**). Male AR100Q mice showed signs of weakness, atrophy and kyphosis and had a median survival of 13 weeks (*χ*^2^ Log-Rank = 17. 1, p < 0.0001) (**Figure 1C**). Female AR100Q mice showed a median survival of 23 weeks (*χ*^2^ Log-Rank = 26.62, p < 0.0001) and presented a phenotype similar to that of male mice at moribundity (**Supplementary Figure 6**). AR100Q mice showed progressive body weight loss starting from 8 weeks of age (p = 0.0001) (**Figure 1D**), and deterioration of motor function assessed by rotarod and hanging wire tasks, starting from week 12 (p = 0.002) and week 11 (p = 0.0001), respectively (**Figure 1E**). AR100Q mice developed atrophy of glycolytic muscles, resulting in 25-to-50% weight loss of quadriceps, gastrocnemius, TA, and EDL, but not soleus, by 8 weeks of age, consistent with previous findings (**Figure 1F**) (14, 30). By nicotinamide adenine dinucleotide (NADH) staining we found a significant (p = 0.0001) glycolytic-to-oxidative fiber-type switch in 8-week-old AR100Q mice, as previously described in SBMA knock-in mice and patients (**Figure 1G**) (26, 30). The median cross sectional area (CSA) of glycolytic and oxidative fibers was decreased by 9% and 8% in 4-week-old AR100Q mice, and by 49% and 11% in 8-week-old mice, respectively, thereby revealing that atrophy of glycolytic fibers exceeded that of oxidative fibers, as previously reported (30). We next analyzed muscle force. By grip strength analysis muscle force was reduced by 38% in 8-week-old AR100Q mice (**Figure 1H**). Altogether, these results show that the newly generated AR100Q mice express physiological levels of hAR and recapitulate the most important features of SBMA, which are deposition of AR in aggregates and inclusion bodies in vulnerable cells, reduced life span, progressive body weight loss, motor dysfunction, and muscle atrophy and weakness, as described in other mice modelling SBMA (11, 12, 32).

**Figure 1.**
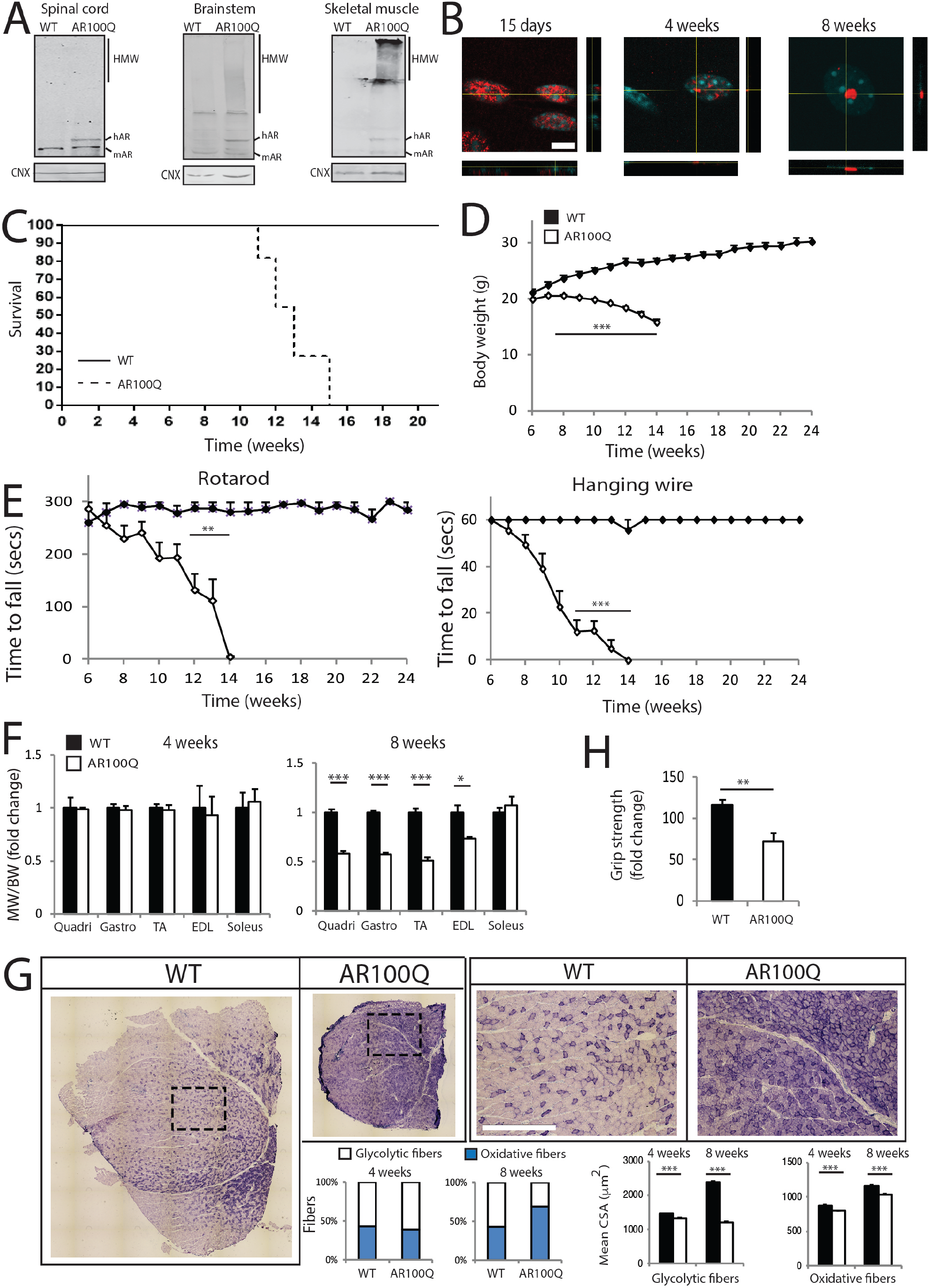
AR100Q transgenic mice show reduced life span and progressive aggregation, motor dysfunction and muscle pathology. A) Western blotting analysis of AR in the spinal cord, brainstem and quadriceps muscle of 8-week-old wild type (WT) and AR100Q mice (n = 3). B) Confocal microscopy analysis of AR subcellular localization in intact myofibers from gastrocnemius of AR100Q mice (n = 3). Bar, 10 micron. C) Kaplan-Meier survival curves compared using Log-rank (Mantel-Cox) test of WT (n = 8) and AR100Q (n = 11) mice. D) Temporal changes in mean body weight (BW) of WT (n = 8) and AR100Q (n = 14) mice. E) Rotarod and hanging wire tasks of WT (n = 8) and AR100Q (n = 14) mice. F) Muscle weight (MW) normalized to BW in WT (n = 14) and AR100Q (n = 16) mice. G) NADH analysis of quadriceps of WT and AR100Q mice (n = 3). Inset position is shown by the dashed square box. Bars, 500 micron. Analysis of the mean CSA of glycolytic and oxidative fibers in the gastrocnemius of WT and AR100Q mice (n = 3). Number of fibers 4-week-old: 1224 WT, and 1237 AR100Q; 8-week-old: 1232 WT, and 1219 AR100Q. H) Grip strength analysis in 8-week-old WT and AR100Q mice (n = 8). Graphs, mean ± SEM, student’s t-test. (A) AR was detected with a specific antibody, and calnexin (CNX) was used as loading control. (B) AR was detected with specific antibody (red), and nuclei with DAPI (blue), and shown are representative images from at least 3 mice for each group. * p < 0.05, ** p < 0.01, *** p < 0.001.

### AR100Q mice develop muscle atrophy and show reduced intrinsic muscle force generation

To question the intrinsic capability of muscle to generate force in AR100Q mice, we performed direct *ex vivo* electrical stimulation, i.e. bypassing neuronal stimulation, of EDL, as a representative fast-glycolytic muscle that shows signs of severe atrophy, and soleus, as a representative slow-oxidative muscle that is less severely affected in SBMA mice (30). Specific peak twitch force (P_t_) and tetanic force (P_o_) production was significantly decreased by about 75% and 71% in the EDL of 8-week-old AR100Q mice (**Figure 2A-B**). Peak twitch force (P_t_) and tetanic force (P_o_) production was decreased by 28% and 37% in soleus, and this difference was not significant, which is consistent with previous findings that slow-twitch muscles are less severely affected in SBMA mice (30). We found an increase in fiber contraction and half-relaxation time, indicating slower contraction kinetics in the EDL muscle of AR100Q mice (**Figure 2C**). Importantly, slower relaxation time also caused a left-shift in the normalized force-frequency curve (**Figure 2D**), showing an anticipated fusion of the stimuli and maximal tension reached at 60Hz for AR100Q EDL muscle compared to 120Hz for the WT EDL muscle at 8 weeks of age. This phenomenon was not observed in soleus. Then we asked whether the abnormalities detected in the EDL muscle occur at 4 weeks of age. We did not find any difference in peak twitch force (P_t_) and tetanic force (P_o_) production in 4-week-old AR100Q mice compared to control mice (**Figure 2E**). Rather, we found that the increased time in fiber contraction and half-relaxation and the shift in the normalized force-frequency curve were already present at 4 weeks of age (**Figure 2F-G**). These findings show that the intrinsic force of fast-twitch muscles is compromised in AR100Q mice and highlight that this aspect of SBMA muscle pathology occurs early in this mouse model of SBMA.

**Figure 2.**
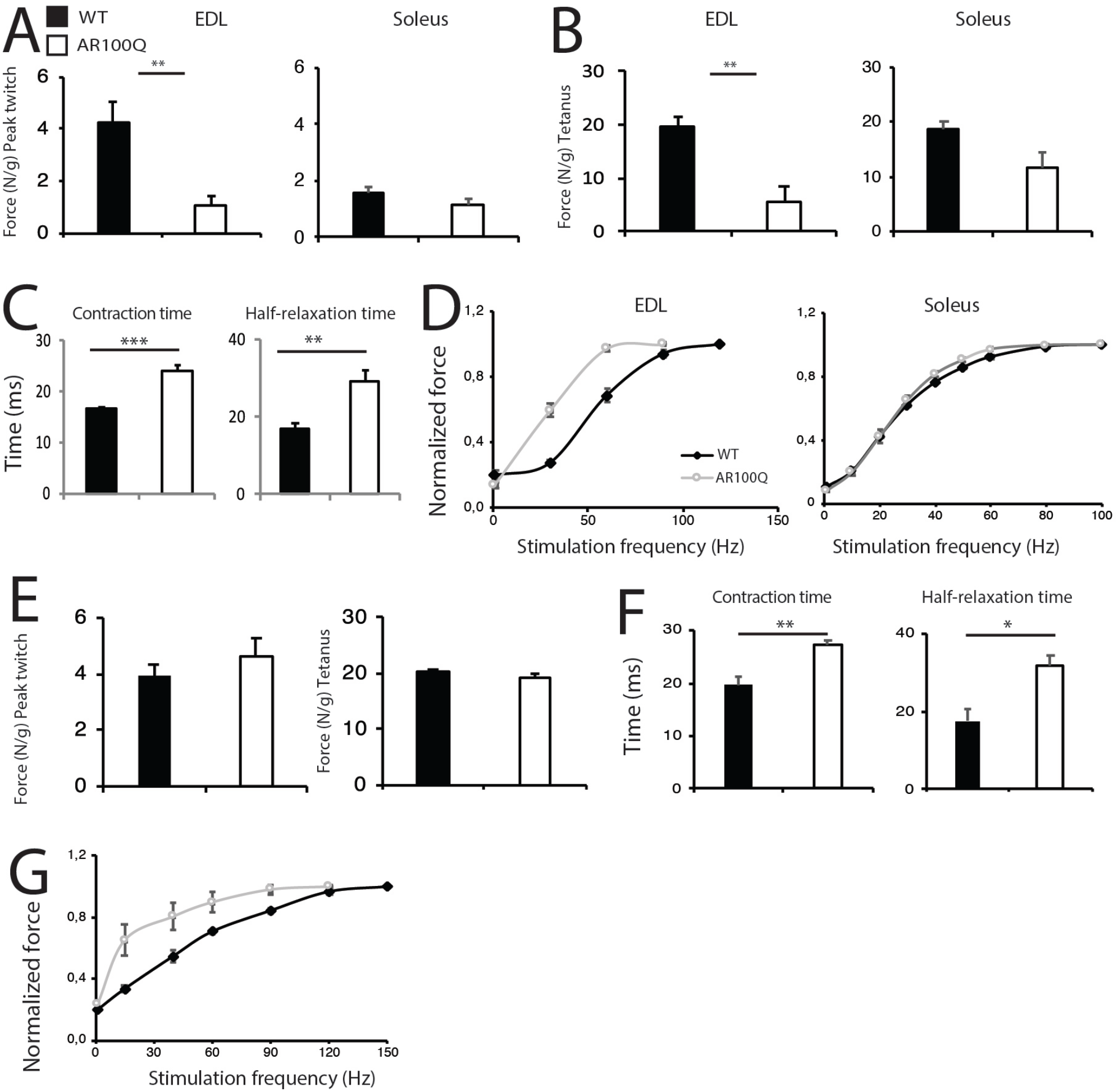
Precocious and progressive alterations in both intrinsic muscle force generation and contraction and half-relaxation time in AR100Q mice. A-B) Intrinsic muscle force generation upon peak twitch (A) and tetanic stimulation (B) in 8-week-old WT and AR100Q mice (n = 4). C) Peak contraction and half-relaxation time in the EDL of 8-week-old WT and AR100Q mice (n = 4). D) Intrinsic muscle force/frequency in 8-week-old WT and AR100Q mice (n = 4). E) Intrinsic muscle force generation upon peak twitch (A) and tetanic stimulation (B) in the EDL of 4-week-old WT and AR100Q mice (n = 4). F) Peak contraction and half-relaxation time in the EDL of 4-week-old WT and AR100Q mice (n = 3). G) Intrinsic muscle force/frequency in the EDL of 4-week-old WT and AR100Q mice (n = 3). Graphs, mean ± SEM, student’s t-test.

### Denervation is a late event in AR100Q mice

To decipher what are the cellular determinants leading to muscle weakness in 100Q mice, we next interrogated motor unit integrity. As previously reported (11, 32, 36), no changes were detected by Nissl staining in the number and soma area of MNs in transversal lumbar spinal cord sections of 12-week-old AR100Q mice compared to WT siblings (**Figure 3A**). By toluidine blue staining of semi-thin sciatic nerve transversal sections, we did not find any gross abnormalities in both the number of axons and g-ratio (axon diameter/nerve diameter) in 8-week-old AR100Q mice, indicating that myelin thickness and axon diameter are preserved (**Figure 3B**). Consistent with previous findings (37), neuromuscular junctions (NMJs) showed a one-to-one matching between acetylcholine receptor (AchR) clusters, detected by α-Bungarotoxin, and the nerve terminals, stained for the presynaptic protein marker syntaxin-1A/1B, in the gastrocnemius, soleus, EDL, TA, and quadriceps (**Figure 3C and data not shown**). Importantly, we found upregulation of denervation markers, including embryonic myosin heavy chain 3 (*Myh3*), perinatal myosin heavy chain 8 (*Myh8*), myogenin (*MyoG*), and muscle-specific kinase (*Musk*), in 8- and 12-week-old, but not 4-week-old AR100Q mice (**Figure 3D**). By hematoxylin and eosin (H/E) staining, we found severe muscle pathology at late stage (12 weeks) of disease characterized by a high grade of fiber size variability, high number of atrophic fibers, often angulated and grouped up to 16-18 fibers, presence of large hypertrophic fibers with central nuclei, which is a feature of SBMA muscle (28), a modest increase of perimysium connective tissue, all together revealing a picture of denervation atrophy with myopathic signs (**Figure 3E**), which recapitulate the muscle pathology described in SBMA patients (27). These observations indicate that denervation is a late event in this mouse model of SBMA, as it is in knock-in SBMA mice (11, 30), and imply that reduced muscle strength precedes denervation, suggesting that it results from early pathological processes mainly ascribable to muscle.

**Figure 3.**
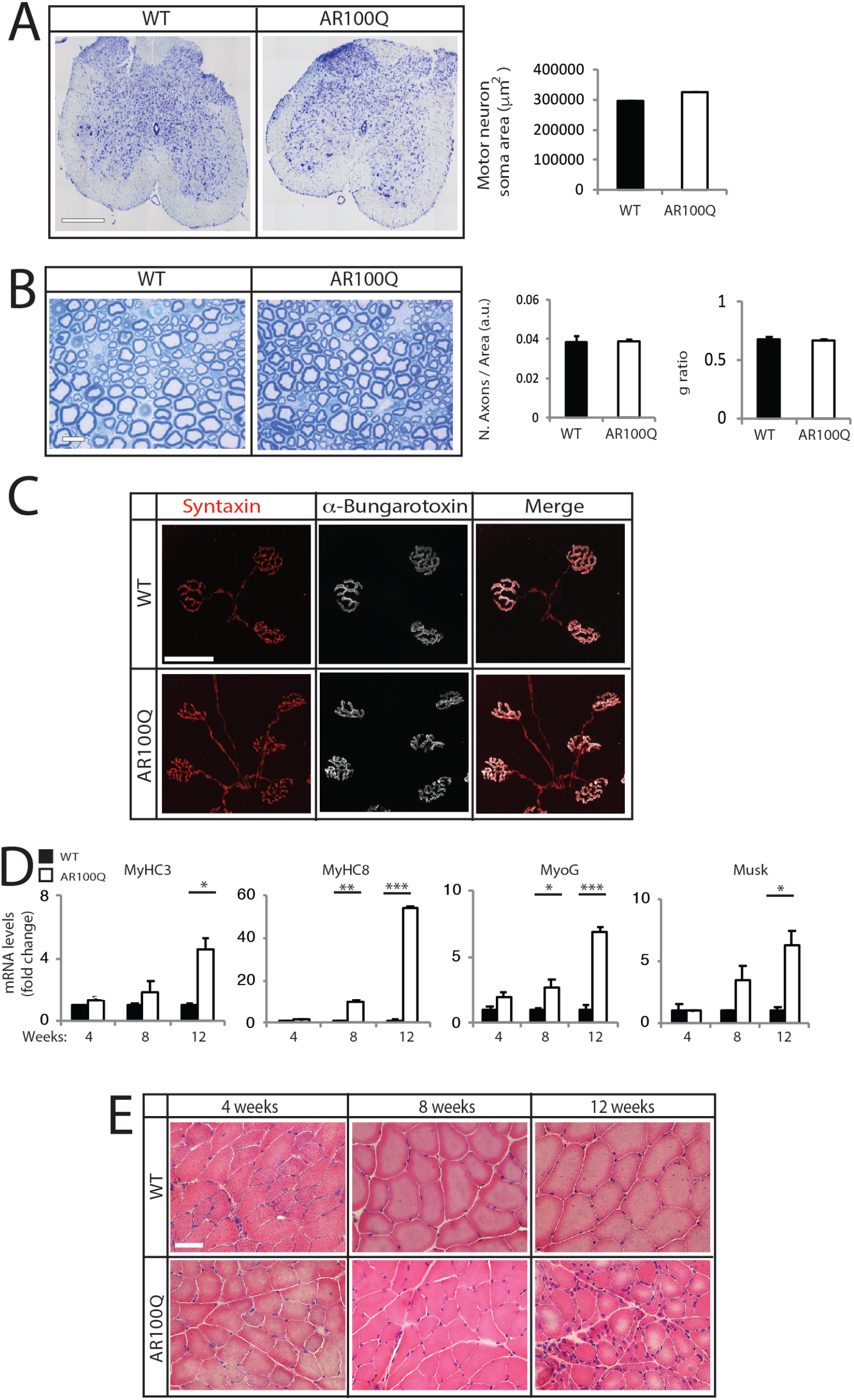
Denervation is a late event in AR100Q mice. A) Nissl staining analysis of motor neuron soma area of lumbar spinal cord transversal sections of 12-week-old WT and AR100Q mice (n = 3). Bar, 500 micron. B) Toluidine blue staining of semi-thin sciatic nerve transversal sections of 8-week-old WT and AR100Q mice (n = 3). Bar, 10 micron. C) Immunohistochemical analysis of NMJ pathology in the quadriceps muscle of 8-week-old WT and AR100Q mice (n = 3). Syntaxin was detected with a specific antibody, and nicotinic acetylcholine receptors with fluorescently labelled a-Bungarotoxin. Bar, 50 micron. D) Real-time PCR analysis in the quadriceps of WT and AR100Q mice (n = 3-5). E) Hematoxylin and eosin staining of transversal sections of quadriceps muscle of WT and AR100Q mice. Bar, 50 micron. Shown are representative images. Graphs, mean ± SEM, student’s t-test (n = 3-5); * p < 0.05, ** p < 0.01, *** p < 0.001.

### AR100Q mice display early fast-to-slow fiber-type switch and altered muscle striation

We next asked whether altered contraction kinetics and reduced force generation are associated with a change in fiber-type composition and muscle triad structural organization. By immunofluorescence analysis of myosin heavy chain (MyHC) subtypes, we found that fiber-type I, IIa, and IIx fibers were increased by 2.5%, 7%, and 11.3%, respectively, whereas type IIb fibers were decreased by 26.4% in 8-week-old AR100Q mice compared to WT mice (**Figure 4A**). Importantly, these fiber-type changes were detected as early as 4 weeks of age. We next analyzed skeletal muscle striation by staining FDB-isolated fibers with an antibody recognizing ryanodine receptor (RYR) 1, the main RYR isoform expressed in skeletal muscle cells. RYR detects the position of Ca^++^ release units in the sarcoplasmic reticulum (SR) and is positioned at the junctional SR (J-SR), which is the region of the terminal SR cisternae that faces transverse tubules (t-tubules) at triads in between longitudinal SR (L-SR) (**Figure 4B**). By confocal microscopy and fluorescence intensity profile, we found that RYR1 formed a regular doublet pattern in WT mice, whereas the organization of Ca^++^ release units was altered in 4- and 8-week-old AR100Q mice (**Figure 4C-D**). These observations indicate that polyQ-expanded AR causes an early and progressive shift from type IIb-to-type IIx and IIa fibers with concurrent alteration of muscle triad organization, likely contributing to the decrease in intrinsic muscle force generation observed before denervation.

**Figure 4.**
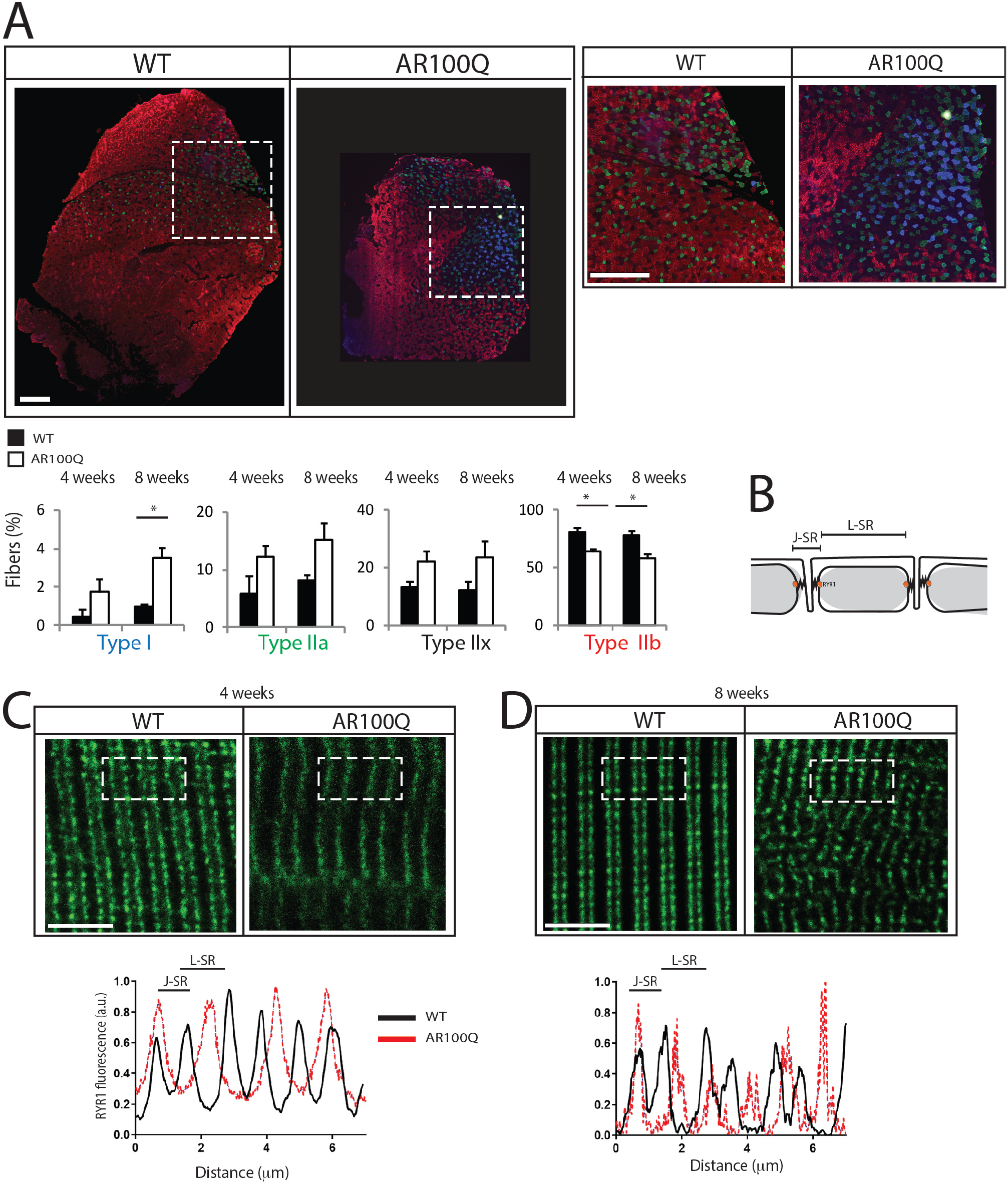
Switch from type IIb to type IIa and IIx fibers and altered muscle striation in AR100Q mice. A) Immunofluorescence analysis of myosin heavy chain (MyHC) type I (blue), IIa (green), IIx (black), and IIb (red) of WT and AR100Q mice. Shown are representative images from 8-week-old mice. Inset position is shown by the dashed square box. Bars, 5 micron. Graphs, mean ± SEM, n = 3-5, student’s t-test. * p < 0.05. B) Schematic representation of the triad. Junctional SR, J-SR; longitudinal SR, L-SR. C-D) Immunofluorescence analysis of RYR1 (green) in FDB-isolated fibers from WT and AR100Q mice (n = 3). Dashed square box indicates the area considered for fluorescence intensity analysis. Bars, 500 micron.

### Activation of catabolic pathways and mitochondrial pathology are late events in the muscle of AR100Q mice

By immunofluorescence of transversal sections of the quadriceps muscle we found reduced RYR1 immunoreactivity in AR100Q mice (**Figure 5A and Supplementary Figure 7**). Notably, in some fibers we noticed areas deprived of RYR1 staining, which resemble the muscle pathology that we previously described in SBMA patients (28). These abnormalities were exclusively found in fast-fibers and were not detected at 4 weeks of age. Alterations in the RYR1 function and muscle striation are associated with processes that lead to mitochondrial dysfunction and activation of catabolic pathways and autophagy (38). We found increased transcript levels of genes involved in protein degradation via the ubiquitin-proteasome system (UPS) and autophagy, such as the E3 ubiquitin ligases, specific of muscle atrophy and regulated by transcription (*Smart*) and muscle ubiquitin ligase of the SCF complex in atrophy-1 (*Musa1*), forkhead box O3a (*Foxo3a*), microtubule-associated protein 1A/1B-light chain 3 (*LC3*), and sequestosome 1 (*Sqstm1*, p62), specifically at 8, and not 4, weeks of age, suggesting increased UPS and autophagy activity (**Figure 5B**). By Western blotting, we found accumulation of LC3II, suggesting increased number of autophagosomes, and p62, suggesting an induction of autophagy, as previously reported in other SBMA mouse models and patients (**Figure 5C**) (30, 39, 40). By measuring mitochondrial membrane potential in myofibers isolated from the FDB muscle upon treatment with the F_1_F_0_-ATPase blocker, oligomycin, we found that the number of fibers with mitochondria depolarized by oligomycin was significantly (P < 0.05) increased in 8-, and not 4-week-old AR100Q mice (**Figure 5D**). By realtime PCR analysis, peroxisome proliferator-activated receptor gamma coactivator 1-alpha (*Ppargc1a*, PGC1α), which promotes mitochondrial biogenesis (41) and whose overexpression shifts muscle metabolism towards oxidative phosphorylation (42), and the complex II respiratory chain subunit, succinate dehydrogenase [ubiquinone] flavoprotein subunit (*Sdha*), were upregulated in the quadriceps of 8-week-old, but not 4-week-old, AR100Q mice (**Figure 5E**). Mitochondrial depolarization and dysfunction are often associated with oxidative stress and upregulation of genes involved in the oxidative stress response, such as nuclear factor erythroid 2 like 1 (*Nfe2l1*, NRF1), and nuclear factor erythroid 2 like 2 (*Nfe2l2*, NRF2). NRF1 and NRF2 are negatively regulated by kelch-like ECH-associated protein 1 (*keap1*) and induce the expression of target genes involved in the oxidative stress response, such as NADPH quinone oxidoreductase (*Nqo-1*) (43). The transcript levels of *Nfe2l1* and *Nqo-1* were upregulated, whereas those of *keap1* were downregulated in the muscle of 8-week-old AR100Q mice (**Figure 5F**). These results indicate that mitochondrial pathology and activation of protein degradation pathways are late events in SBMA muscle.

**Figure 5.**
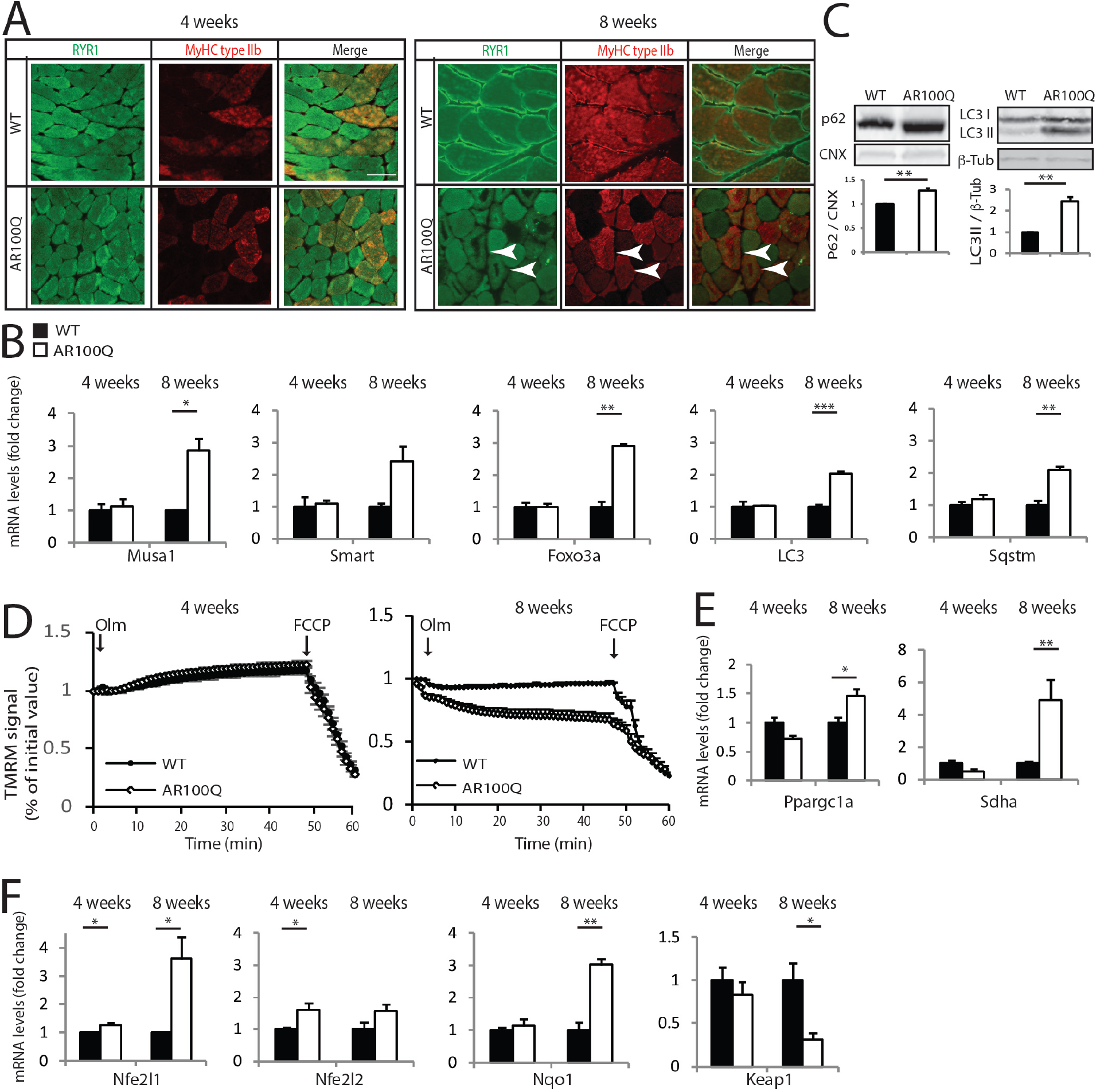
Mitochondrial pathology and activation of catabolic pathways are late events in the fast-twitch muscles of AR100Q mice. A) Immunofluorescence analysis of RYR1 (green) and MyHC type IIb (red) in the quadriceps of WT and AR100Q mice (n = 3). Shown are representative images. Bar, 50 micron. Arrows indicate fibers with core-like pathology. B) Real-time PCR analysis in the quadriceps of WT and AR100Q mice (n = 3-5). C) Western blotting analysis in 8-week-old WT and AR100Q mice (n = 3). D) Mitochondrial membrane depolarization in FDB-isolated fibers of WT and AR100Q mice (n = 3). Olm, oligomycin; FCCP, carbonyl cyanide-4-(trifluoromethoxy)phenylhydrazone. E-F) Real-time PCR analysis in the quadriceps of WT and AR100Q mice (n = 3-5). Graphs, mean ± sem, student’s t-test; * p < 0.05, ** p < 0.01, *** p < 0.001.

### Expression of polyQ-expanded AR alters Ca^++^ homeostasis and affects the expression of ECC genes

A key factor that affects muscle force generation is the alteration of Ca^++^ dynamics during muscle contraction. Muscle contraction and force generation elicited by MN impulse trigger a process known as the ECC, which links the electrical events occurring in the sarcolemma with myoplasmic Ca^++^ dynamics to allow myofiber contraction (44). Upon arrival of the action potential at the NMJ, membrane depolarization leads to a conformational change in dihydropyridine receptors (DHPRs) and RYR, leading to release of Ca^++^ from the SR to the myoplasm to trigger muscle contraction. Ca^++^ transients in the myoplasm are rapidly followed by Ca^++^ reuptake through the SR Ca^++^-ATPase (SERCA) 1 and SERCA2, to stop contraction and relax myofibers (45). The magnitude of Ca^++^ transients is regulated by several buffering proteins, including parvalbumin (PV) in the cytosol (46), and calsequestrin 1 (CASQ1) and CASQ2 in the SR (47). Another key player in Ca^++^ dynamics is the SERCA inhibitor, sarcolipin (SLN) (45). Our observation that time to contract and relax was extended in AR100Q mice (**Figure 2C and F**) prompted us to test whether aberrant handling of Ca^++^ dynamics contributes to defects in intrinsic muscle force generation. We subjected enzymatically-dissociated FDB fibers loaded with the ratiometric Ca^++^ dye Fura-2 to both isolated twitches and prolonged high-frequency stimulation (60 Hz, 2s). Upon single twitch stimulation, the kinetics to reach the peak in cytosolic Ca^++^ was prolonged in AR100Q mice (**Figure 6A**), suggesting impaired balance between Ca^++^ release from SR stores to the cytosol and Ca^++^ reuptake through SERCA. Moreover, we found a significant delay in the decrease of Fura-2 signal in AR100Q mice, suggesting a slow kinetics of myoplasmic Ca^++^ clearance (**Figure 6B**). Similar effects were observed upon tetanic stimulations, where a slower cytosolic Ca^++^ depletion was observed in AR100Q mice (67.1 ms in AR100Q vs 55.4 ms in WT mice) (**Figure 6C**). However, the slower depletion was not accompanied by a statistically significant increase in the total cytosolic Ca^++^ (**Supplementary Figure 8A**), because the impaired balance of Ca^++^ release, reuptake and buffering was maintained also in fast transients between stimuli, leading to a smaller increment for each oscillation (ΔR = 0.08 vs. 0.12 in WT mice, p < 0.0001) (**Figure 6D and Supplementary Figure 8B**). These results suggest that during intense myofiber activity transgenic mice have impaired Ca^++^ dynamics across the different subcellular compartments.

**Figure 6.**
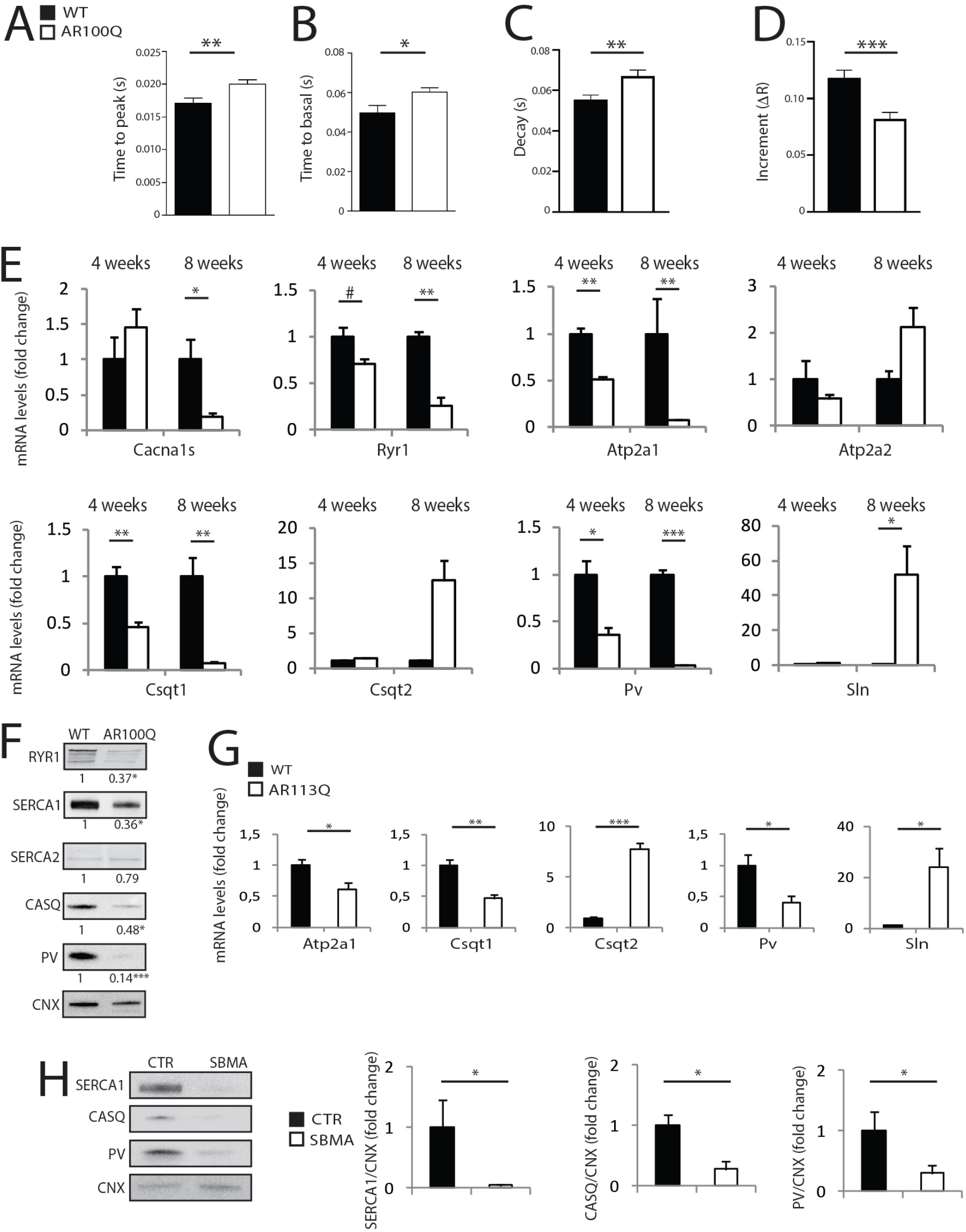
Altered Ca^++^ dynamics associated with ECC gene expression changes in SBMA muscle. A-D) Analysis of Ca^++^ levels in response to twitch (A-B) and tetanic (C-D) stimulation in FDB-isolated fibers of 8-week-old WT and AR100Q mice (n = 3). E) Real-time PCR analysis in the quadriceps of WT and AR100Q mice (n = 3-5). F) Western blotting analysis in 8-week-old WT and AR100Q mice (n = 3). Quantification is shown at the bottom of each panel. G) Real-time PCR analysis in the quadriceps of 26-week-old WT and AR113Q mice (n = 3-5). H) Western blotting analysis in quadriceps muscle biopsies obtained from SBMA patients and control (CTR) subjects (n = 3-6). Graphs, mean ± sem, student’s t-test; # p = 0.059, * p < 0.05, ** p < 0.01, *** p < 0.001.

In a microarray previously performed in quadriceps muscle of SBMA knock-in mice (30), the most upregulated gene was *Sln*. By real-time PCR, we found that the mRNA transcript levels of key ECC genes, including *Cacna1s*, which encodes a subunit of the DHPR, *Ryr1, Atp2a1, Csqt1*, and *Pv*, were significantly decreased, whereas those of *Atp2a2, Csqt2*, and *Sln* were upregulated in the quadriceps, EDL, and FDB muscles of 8-week-old AR100Q mice (**Figure 6E and Supplementary Figure 8C-D**). These gene expression abnormalities were attenuated or absent in the soleus muscle, indicating that these alterations mainly occur in fast-glycolytic muscles (**Supplementary Figure 8E**). Importantly, changes in the expression of *Ryr1, Atp2a1, Csqt1, Csqt2, Pv*, and *Sln* were detected as early as 4 weeks of age at both mRNA and protein level in AR100Q mice (**Figure 6E-F**), as well as knock-in AR113Q mice (**Figure 6G**). Notably, these alterations were also present in muscle biopsy specimens derived from SBMA patients (**Figure 6H**). These results suggest that changes in the expression of ECC genes detected in two different SBMA mouse models and in SBMA patients and may affect myoplasmic Ca^++^ clearance, which in turn explains the altered dynamics of muscle contraction found in our transgenic mice, as shown here, and in SBMA patients (48).

### Alteration in ECC gene expression is AR signaling-dependent and can be pharmacologically reverted *in vivo*

The results described above prompted us to establish whether expression of ECC genes is modified upon acute modulation (enhancement and suppression) of AR expression and transactivation in the adulthood. To acutely induce polyQ-expanded AR expression *in vivo*, we generated transgenic mice for doxycycline (dox)-inducible expression of hAR with 100 glutamine residues (hereafter referred to as iAR100Q for inducibleAR100Q), which were subsequently crossed with mice expressing rtTA under the CMV promoter (hereafter referred to as rtTA mice) (49). iAR100Q /rtTA double transgenic mice were fertile, viable and did not show any overt phenotype in the absence of dox. Administration of dox (0.1, 0.2, and 1g/L) to pregnant dams did not result into double-positive pups, indicating that induction of *hAR* trangene is highly toxic for embryo development, possibly due to high copy number insertion (**Supplementary Figure 9 and data not shown**). To induce polyQ-expanded AR expression in adulthood, thereby avoiding compensatory or toxic effects occurring during development and before sexual maturation, we started dox treatment at 6 weeks of age (**Figure 7A**), which more or less corresponds to disease onset in transgenic SBMA mice constitutively expressing mutant AR (**Figure 1**). While treatment of iAR100Q/rtTA mice with 0.1 and 0.2 g/L of dox was not sufficient to induce hAR expression and trigger disease manifestations, administration of 1 g/L of dox efficiently induced hAR expression (**Figure 7B**), ultimately causing premature death (p = 0.001) with a median survival of 8 weeks of age (maximum life duration was 15 weeks of age) (**Figure 7C**). Similar to the transgenic mice constitutively expressing mutant AR, iAR100Q/rtTA mice manifested a progressive loss of body weight and reduced muscle strength (**Figure 7D-E**), a phenotype not observed in WT, rtTA, and iAR100Q mice (**Supplementary Figure 10A-D**). Dox treatment reduced quadriceps, and not soleus, muscle mass by 21% and CSA by 52% of iAR100Q /rtTA mice (**Figure 7F-G**), but not in WT, rtTA, and iAR100Q mice (**Supplementary Figure 10E**). Importantly, these observations indicate that post-natal induction of AR100Q expression is sufficient to cause muscle atrophy and to induce a phenotype that recapitulates the main features of SBMA. The transcript levels of *Atp2a1* did not change, whereas those of *Csqt1, Csqt2*, and *Pv* were significantly reduced and those of *Sln* were significantly increased in iAR100Q/rtTA mice in response to dox treatment (**Figure 7H**). These results indicate that acute induction of expression of polyQ-expanded AR in the adulthood is sufficient to induce changes in the expression of key genes involved in muscle contraction and relaxation similar to those observed in the transgenic mice constitutively expressing AR100Q.

**Figure 7.**
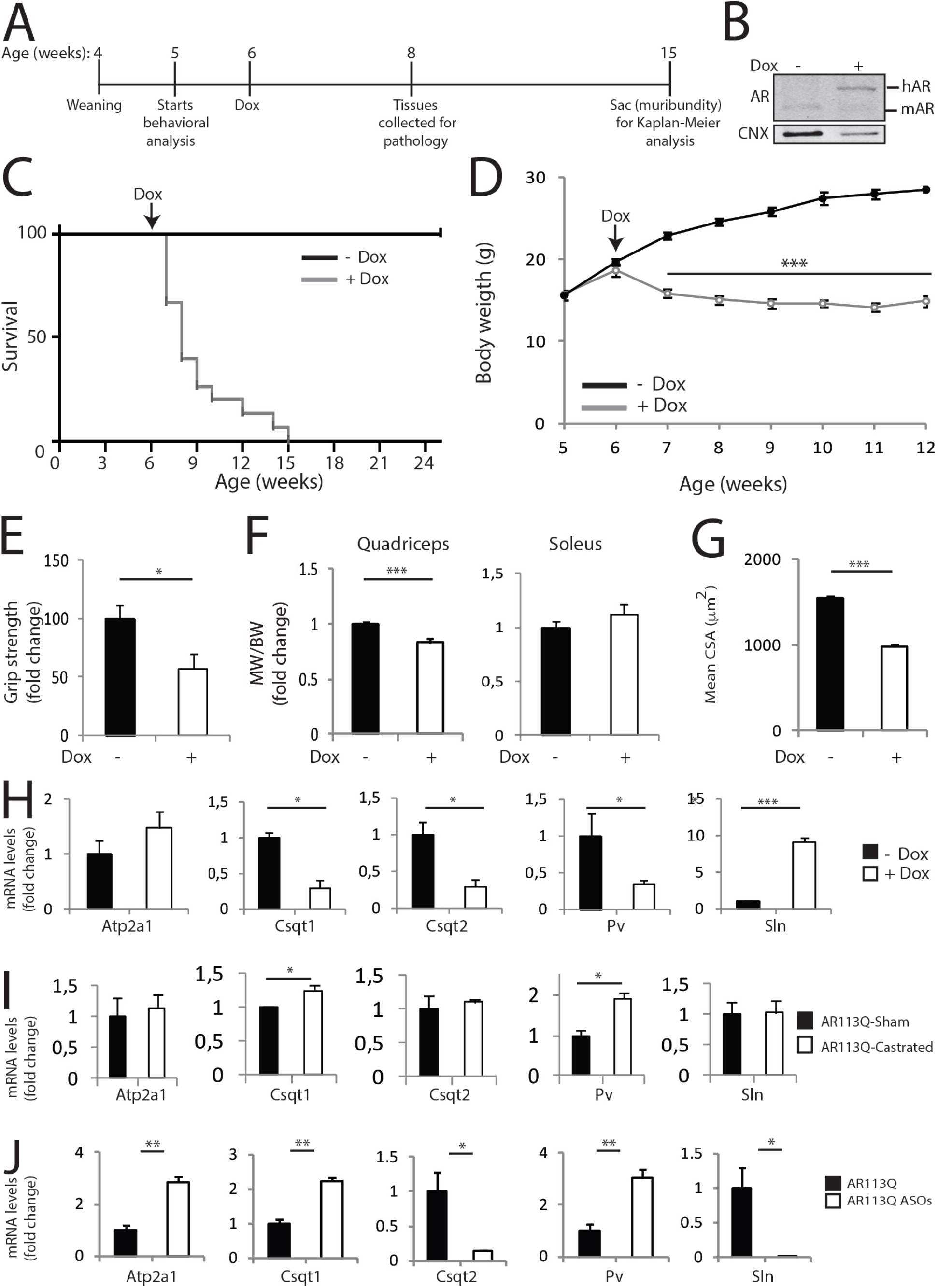
Modulation (induction and suppression) of polyQ-expanded AR expression in the adulthood modifies ECC gene expression. A) Scheme of treatment of conditional SBMA mice with Dox. B) Western blotting analysis of AR expression in the quadriceps of iAR100Q /rtTA mice treated with either vehicle (-Dox) or dox (+ Dox). Shown is one experiment representative of three. C) Kaplan-Meier analysis of survival of iAR100Q /rtTA mice treated with either vehicle (n = 8) or dox (n = 15). Survival curves were compared using Log-rank (Mantel-Cox) test. D) Temporal changes in mean BW of iAR100Q /rtTA mice treated with either vehicle (n = 8) or dox (n = 15). E) Grip strength analysis of muscle force of iAR100Q /rtTA mice treated with either vehicle (n = 8) or dox (n = 8). F) MW normalized to BW in iAR100Q /rtTA mice treated with either vehicle (n = 4) or dox (n = 5). G) Analysis of the mean CSA of fibers in the quadriceps of iAR100Q /rtTA mice (n = 3) treated with either vehicle or dox. Number of fibers: 4400 vehicle, 4100 dox. H-J) Real-time PCR analysis in the (H) quadriceps of iAR100Q /rtTA mice (n = 3-5) treated with either vehicle or dox, (I) levator ani/bulbocarnosus (LA/BC) muscle of 26-week-old AR113Q mice (n = 3-5), and (J) quadriceps muscle of 26-week-old AR113Q mice treated with either vehicle or ASOs (n = 3-5). Graphs, mean ± sem, student’s t-test; * p < 0.05, ** p < 0.01, *** p < 0.001.

To inhibit polyQ-expanded AR *in vivo*, we undertook two different approaches. To prevent polyQ-expanded AR transactivation by androgens we performed surgical castration, and to suppress its expression we treated the mice with ASOs. Both approaches had beneficial effects on SBMA phenotype (10, 11, 20, 22, 34). Surgical castration increased the expression of *Csqt1* and *Pv* in AR113Q mice (**Figure 7I**), whereas suppression of polyQ-AR expression by ASOs treatment restored the expression of the all ECC genes (**Figure 7J**). These results indicate that acute modulation (induction and suppression) of AR expression modifies the expression of key genes regulating muscle contraction and Ca^++^ dynamics and suggest that these gene expression changes are valuable pharmacologic targets for SBMA patients.

### Direct AR regulation of ECC gene expression is altered in SBMA muscle

The native function of polyQ-expanded AR is a key component of disease pathogenesis (50, 51). AR is a transcription factor activated by androgens, whose function is altered upon polyQ expansions. We searched for androgen-responsive elements (AREs) in the promoters of the ECC genes whose expression is altered in SBMA muscle. By a computational biology approach, we found several AREs located in the enhancers, promoters, and 5’-untranslated regions (5’-UTR) of these genes, suggesting a direct involvement of AR in their transcription (**Figure 8A and Supplementary Table 1**). We reasoned that if these genes are under the control of androgen signaling in physiological conditions, their expression is predicted to change when the androgen-induced AR transactivation is inhibited. Consistent with this idea, suppression of androgen-dependent AR transactivation by surgical castration in WT mice significantly decreased the expression of *Csqt1* and *Pv*, suggesting that these genes are positively regulated by AR, and it increased the expression of *Sln*, suggesting that this gene is negatively regulated by AR in physiological conditions (**Figure 8B**). Importantly, these gene expression changes resembled those observed in SBMA muscle, suggesting that polyQ expansion affects AR native function in pathological conditions. To further explore this possibility, we focused on *sln* that was the most up-regulated gene among the differentially expressed genes (DEGs) identified in SBMA muscle (30). To determine whether AR binds to *sln* promoter, we performed chromatin immunoprecipitation (ChIP) of AR in the muscle of WT and AR100Q mice. We found that AR binds to the *sln* promoter in physiological conditions, and AR promoter occupancy was lost in AR100Q mice (**Figure 8C**). These results suggest that a novel native physiological function of AR, i.e. androgen-dependent regulation of expression of ECC genes, is disrupted in pathological conditions.

**Figure 8.**
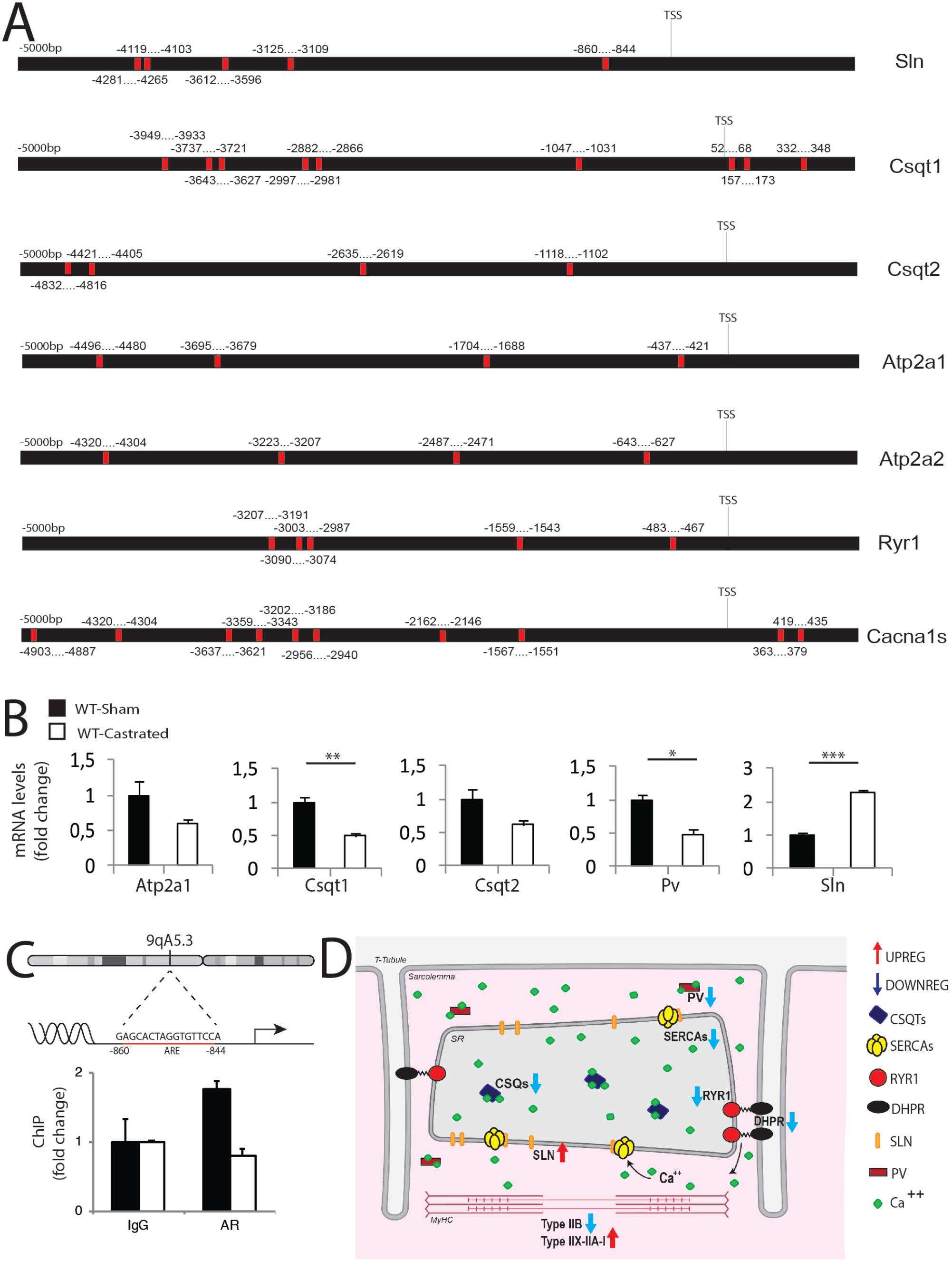
Androgen-dependent AR regulation of ECC gene expression is disrupted in SBMA. A) Bioinformatic analysis of AREs upstream and downstream the transcription start site (TSS, +1) of the indicated ECC genes in mouse. B) Real-time PCR analysis in the LA/BC muscle of 26-week-old WT mice (n = 3-5). C) ChIP analysis of AR occupancy at the indicated ARE on the promoter of the *Sln* gene. D) Schematic representation of ECC gene expression changes occurring in SBMA muscle. Graphs, mean ± sem, student’s t-test; * p < 0.05, ** p < 0.01, *** p < 0.001.

These findings implicate a role for androgen signaling in the regulation of muscle contraction and Ca^++^ homeostasis in physiological conditions, a level of regulation of gene expression in muscle that is perturbed in SBMA and may play a role in pathologies involving loss of androgen signaling (**Figure 8D**).

## Discussion

Although skeletal muscle is considered nowadays as a primary site of toxicity in SBMA, the mechanisms through which polyQ expansions in the AR alter muscle mass, function and homeostasis is poorly understood. Using two novel SBMA mouse models for constitutive and inducible expression of the disease protein, together with knock-in mice and human muscle biopsy samples, here we provide the first evidence that mutant AR alters the pathways representing the groundwork for muscle function (fiber-type myosin), structure (triad), and Ca^++^ dynamics. These events occurred before denervation and involved alteration of the native function of AR.

A unifying symptom in SBMA patients is a slow, though progressive, loss of muscle strength, estimated around 2% per year (22, 24). This feature is invariably recapitulated in all the animal models of SBMA, including those used here. Moreover, force decay is accompanied by a profound alteration in the kinetics of muscle contractility (29, 52). However, a clear mechanism explaining these alterations and how they affect force in SBMA muscle was still missing. We found a profound rearrangement in the expression of MyHC isoforms, with SBMA muscles displaying a generalized “slowing” of myosin types. In terms of contraction strength, the difference in power released between slow and fast MyHC can be up to 10-fold (53). Thus, the switch in MyHC types may contribute to the loss of muscle force and performance of SBMA muscles. Moreover, the muscle of AR100Q mice was characterized by abnormal muscle striation, with progressive altered organization of the Ca^++^-release units in AR100Q muscles. Fiber-type change and alteration of muscle striation occurred before denervation, suggesting that they result from intrinsic pathogenetic processes occurring in muscle.

An unprecedented key finding here is that SBMA muscle is characterized by altered Ca^++^ dynamics associated with changes in the expression of ECC genes that regulate muscle contraction and Ca^++^ transients. We demonstrated that SBMA myofibers display a slower release of Ca^++^ within the cytosol upon stimulation and a slower kinetics of myoplasmic Ca^++^ clearance to stop muscle contraction. These findings provide a mechanistic explanation both for the alterations in contractile kinetics of SBMA muscles, and for the significant loss of performance characterizing SBMA muscles. Indeed, disruption and alteration of the ECC machinery at any level result in a range of myopathic conditions, spanning from myotonia to weakness, paralysis and muscle wasting, which are classified as triadopathies (54). Mutations in *CACNA1S* and *RYR* genes lead to malignant hyperthermia and specific forms of congenital myopathy, such as central core disease (55, 56), which are characterized by muscle weakness, centrally located cores, and lack of mitochondria and oxidative enzymes with disorganized contractile apparatus (57). We have previously described a pattern of muscle pathology in SBMA patients that resembles central core disease, with muscle fibers presenting central cores deprived of mitochondria and oxidative enzyme activity (28). Importantly, a similar muscle pathology was detected also in AR100Q transgenic mice. RYR1-related myopathies are often associated with reduced expression and LOF of RYR1, which leads to impaired Ca^++^ release and ECC and reduced muscle force generation. Decreased levels of DHPR, biochemical alteration of RYR and DHPR-RYR1 uncoupling are associated with the intrinsic loss of muscle force in aged muscles (58, 59). We found decreased expression of *Cacna1s* and *Ryr1* in AR100Q. These gene expression changes occurred at late stage of disease, therefore suggesting secondary alterations due to pathological processes occurring in muscle. Nonetheless, reduced expression of *Ryr1* may contribute to the increased time to Ca^++^ release and slower contraction in response to electrical stimulation observed in our transgenic mice. Mutations in the *ATP2A1* gene, leading to increased degradation or decreased activity of SERCA1, are associated with Brody myopathy, a disorder characterized by muscle weakness, delayed muscle relaxation and stiffness due to impaired SR Ca^++^ reuptake. Alteration of SERCA function has been linked also to myotonic dystrophy and hypothyroid myopathy, and decreased activity/concentration of this protein is associated with selective atrophy of fast myofibers (60). Interestingly, myotonia-like symptoms have been reported also in SBMA patients (61), and SERCA1 was decreased in AR100Q mice. Notably, changes in *Atp2a1* expression were detected very early, before motor dysfunction, suggesting that they may play a pathogenic role in SBMA muscle. SERCA activity is regulated by SLN. Increased expression of SLN reduced muscle force, slowed down the rate of contraction and relaxation and altered Ca^++^ reuptake into the SR (62), whereas its ablation had the opposite effect (63). SERCA2 and SLN are upregulated in dystrophic muscles of Duchenne muscular dystrophy (DMD) mice (64), and a gene-therapy approach to decrease SLN levels has been shown to have beneficial effects in DMD mice (65), indicating that increased SLN levels contribute to muscle dysfunction in DMD. SLN expression was also augmented in rodent models of dysferlinopathies and nemaline myopathy (66, 67), and in SBMA, as shown here, where it may contribute to the slow kinetics of Ca^++^ reuptake after muscle contraction.

In addition to SLN, Ca^++^ dynamics are tightly controlled by PV and CASQ1/2. Ablation of CASQ1 altered muscle contractility and increased susceptibility to spontaneous mortality and heat/anesthesia-induced malignant hyperthermia-like episodes (47, 68). Patients with mutations in CASQ1 present with a late-onset exordium with slow progression, and symptoms include high levels of serum CK, fibrofatty substitution, exercise intolerance, mild and age-dependent muscle weakness and atrophy, and the presence of vacuoles reactive to SR Ca^++^-regulating proteins (69). Several of these symptoms are found in SBMA patients (27, 70–72), and deleterious effects of chronic low-intensity endurance exercise were reported in SBMA mice (14). Our findings that CASQ1 expression is regulated by AR signaling and is reduced before onset of atrophy and motor dysfunction in SBMA mice suggest a role for this key ECC gene in pathological processes occurring in SBMA. Consistent with this idea, incidence of lethal hyperthermia is higher in male compared to female CASQ1 null mice, further implying a pathogenic link among the ECC process, sex steroid hormones and their receptors (73). SBMA muscles were almost completely depleted of PV. Reduced PV expression was reported in the arrested development of righting response (*adr*) mouse, an animal model of ADR myotonia characterized by abnormally prolonged muscle contraction and half-relaxation time (74), both present in SBMA fast muscles. Notably, loss of PV expression results in a fast-to-slow fiber-type switch (75), suggesting that PV loss in SBMA muscle may contribute to both fiber-type change and increased rate of relaxation.

In SBMA, mechanisms of AR LOF and toxic GOF coexist and are associated with transcriptional dysregulation, potentially resulting in activation of different transcriptional programs in different subcellular populations. The finding that upon castration wild type mice display a dysregulation in the expression of ECC genes similar to that observed in SBMA muscles suggests that key genes involved in muscle contraction are under the control of androgen signaling. *Sln* was the most upregulated gene in SBMA muscle, its expression was upregulated by surgical castration in normal mice, and it was restored back to normal levels in SBMA mice upon suppression of polyQ-expanded AR toxicity, highlighting this gene as an example of an AR-regulated gene in muscle. Consistent with the idea that *sln* is suppressed by AR, we found that AR occupancy at the *sln* promoter in muscle in physiological conditions. Importantly, AR occupancy was lost in SBMA muscle. The observation that disruption of ECC genes occurs very early in AR100Q mice, even before the manifestation of prominent muscular deficits, suggests that it may be a pathogenic cause of the disease rather than a consequence. Another major result we obtained here is the finding that the alterations in the expression of ECC genes linked to polyQ AR appear to be reversible. Pharmacologic treatment to attenuate the toxicity of polyQ-expanded AR *in vivo* restored the expression of ECC genes. This evidence shows that intervention to attenuate the degeneration induced by polyQ-expanded AR has a positive outcome on gene expression changes associated with pathological processes occurring in muscle, thereby implying a pathogenic role for such gene expression changes in SBMA muscle.

In conclusion, our results identify the ECC machinery as a novel target for developing new therapeutic strategies to attenuate SBMA. Our findings add insights onto the pathological processes occurring in skeletal muscles. Most importantly, our data imply a novel physiological function for AR in controlling the expression of ECC genes, possibly providing a molecular explanation about how AR and its natural ligands carry out their anabolic activity in muscle.

## Materials and methods

### Animals

Animal care and experimental procedures were conducted in accordance with the Italian Institute of Technology, the University of Trento, the University of Padova, and the University of Michigan ethics committees, and were approved by the Italian Ministry of Health. The mice were housed in filtered cages in a temperature-controlled room with a 12-hour light/dark cycle with *ad libitum* access to water and food. By random insertion transgenic mice were generated to express full-length hAR with a pathogenic (AR100Q) polyQ tract. For constitutive AR expression, hAR transgene expression was driven by the cytomegalovirus immediate-early enhancer and the chicken beta-actin (pCAGGS) promoter. For conditional AR expression, hAR transgene expression was driven by the tetracycline-responsive element (TRE). Mice were genotyped by PCR on tail DNA using REDExtract-N-Amp Tissue PCR kit (Sigma-Aldrich) according to manufacturer’s instructions, using the following primers: AR100Q: Forward 5’-CTTCTGGCGTGTGACCGGCG, reverse 5’-TGAGCTTGGCTGAATCTTCC. iAR100Q: Forward 5’-CGTATGTCGAGGTAGGCGTG, reverse 5’-TGAGCTTGGCTGAATCTTCC.Transgenic and inducible lines were backcrossed to the C57Bl6J background for more than 10 generations before subsequent analysis of phenotype and pathology. rtTA mice (Stock n.: 003273) were purchased by The Jackson Laboratory. Generation and genotyping of knock-in AR113Q mice were previously described (11). Mice were treated with doxycycline hyclate (Sigma Aldrich; D9891) in drinking water containing 5% of sucrose (Sigma-Adrich S8501). ASOs treatment and surgical castration were previously described (34). For rotarod, hanging wire and grip strength analyses mice were randomized and the operator was blind for genotype. *ex vivo* muscle force of the gastrocnemius was measured as previously described (76). For survival analysis (GraphPad Prism 5), moribundity was set as the time in which the mouse lost 20% of body weight or showed inability to move, dehydration and cachexia.

### Human samples

Anonymized control and patient biopsy samples were obtained from the Neuromuscular Bank of Tissues and DNA samples, Telethon Network of Genetic Biobanks, and the EuroBioBank network. Tissue collection was approved by the local Ethics Committee, and written informed consent was obtained from each patient. All patients who underwent muscle biopsy were clinically affected and showed weakness and/or fasciculation and/or muscle atrophy, as previously described (28). Myopathic changes together with neurogenic atrophy were observed in muscle biopsies.

### Histological analysis

For muscle histology, muscles were collected immediately after euthanasia, flash-frozen in liquid nitrogen and embedded in optimal cutting temperature (OCT) compound (Tissue Tek, Sakura). Cross sections (10 μm thick) were cut with a cryostat (CM1850 UV, Leica Microsystems) and processed for H/E and NADH staining, as previously described (36). Images were taken using an upright epifluorescence microscope (Zeiss Axio Imager M2) equipped with an X-Cite 120Q fluorescence light source and a Zeiss Mrm Color Camera. Multichannel images and mosaics were taken using Zeiss Axio Vision Software (V.4.8.2 SP3). For Nissl staining of motor neurons in brainstem and spinal cord, deeply anesthetized mice were transcardially perfused with 4% paraformaldehyde (PFA), and tissues were collected, post-fixed in PFA at 4°C for 16 hours and then washed abundantly in phosphate buffered saline (PBS). Tissues were cut and mounted on slices. The samples were gradually dehydrated (70, 95, and 100% EtOH), stained with a solution containing 0.1% cresyl violet for 5 min, washed in water, gradually dehydrated (70, 95, and 100% EtOH), cleared in xylene, and mounted in Eukitt (Bio Optica). All the images were acquired with the inverted Zeiss Observer Z1 microscope. Toluidine blue was performed on sciatic nerve fixed in phosphate buffer 2% glutaraldehyde (Polyscience cat.07710) to assess the number of axons and the myelin thickness. Serial 1 μm thick sections were taken from the nerves of three mice for each group. Images were taken with a 100x objective, on a Leica DM5000 microscope. Analysis of the number of axons and the g-ratio was performed using Image J and the G-Ratio Calculator plugin (http://cifweb.unil.ch).

### Biochemical analysis

The mice of the indicated age and genotype were euthanized and the tissues were immediately collected, snap-frozen in liquid nitrogen, and stored at −80°C until further processing. Mouse tissues were pulverized using pestle and mortar and homogenized in RIPA buffer containing protease inhibitors (Sigma), and 2% sodium dodecyl sulfate (SDS) (**Figure 1A** spinal cord and skeletal muscle and **Supplementary Figure 2A**), and 0.1% SDS (**Figure 1A** brainstem) and. Lysates from muscles and liver were homogenized (homogenizer RZR 2052 control, Heidolph) at 600 rpm (20 times), sonicated, and centrifuged at 15,000 rpm for 15 min at room temperature. Other tissue lysates were sonicated and centrifuged at 15,000 rpm for 15 min at room temperature. Protein concentration was measured using the bicinchoninic acid assay method. For Western blotting analysis, equal amounts of protein were separated in 7.5% tris-HCl SDS-PAGE (for AR), 3-8% NuPAGE Tris-Acetate gels for RYR1, and 4-12% NuPAGE (Thermo Scientific) Bis-Tris gels for ECC proteins. Gels were blotted overnight onto 0.45-mm nitrocellulose membranes (Bio-Rad, 162-0115). Filter retardation assay and ubcellular fractionation were carried out as previously described (36). The following primary antibodies were used: AR (H280 sc-13062, 1:1000), tubulin (T7816, 1:5000), P62 (P0067, 1:1000), LC3B (L7543, 1:1000), Calnexin (ADI-SPA-860, 1:5000), RYR1 (# MA3-925, 1:2000), CASQ (VIIID12; 1:3000), SERCA1 (VE121G9, 1:50000), SERCA2 (N19; 1:1000), PV (ab11427, 1:50000). Protein signals were detected using either the Li-Cor Odyssey Infrared Imaging System or the Alliance Q9 Mini chemidoc system (Uvitec, Cambridge) with appropriate secondary antibodies. Quantifications were performed using ImageJ 1.45 software. Mitochondrial depolarization was performed as previously described (30).

### Immunofluorescence analysis

Muscles were isolated from mice of indicated age and genotype and immediately fixed in 4% PFA. Bladder was used as whole mount preparation, whereas gastrocnemius and FDB were further dissected into muscle bundles of about 20 myofibers. Samples were quenched in 50 mM NH_4_Cl for 30 minutes at RT and then saturated for 2 hours in blocking solution (15% vol/vol goat serum, 2% wt/vol BSA, 0.25% wt/vol gelatin, and 0.2% wt/vol glycine in PBS containing 0.5% Triton X-100). Primary antibodies were diluted in the same solution and then added to muscles for at least 48 hours at 4°C under gentle agitation. Anti syntaxin-1A/1B (rabbit polyclonal, diluted 1:200) was used to stain both motor neuron axons and nerve terminals, as previously described (77). Anti-AR (1:200) was from Santa Cruz (H280 sc-13062). Anti-RYR (1:200) was from Thermo Fisher (MA3-925). Muscles were extensively washed with PBS for at least 2 hours and then incubated for additional 2 hours with appropriate secondary antibodies (Life Technologies) supplemented with fluorescently labelled α-bungarotoxin (Life Technologies) to stain nicotinic acetylcholine receptors, and with Hoechst 34580 (Sigma Aldrich) where indicated. After extensive washes with PBS, muscles were then rinsed with deionized water and mounted on a coverslip with mounting solution (Dako S3023) for microscopy examination. J-SR and L-SR distance was measured using ImageJ. Where indicated, immunofluorescence analysis of RYR1 and AR were performed in 10 μm quadriceps cryosections using the same antibodies listed above supplemented with fluorescent WGA (Thermo Scientific W11261, 1:500). Images were collected with an epifluorescence microscope (Leica CTR6000) equipped with X5 N PL AN 0.12, X20 N PL AN 0.40 objectives or by a SP5 confocal microscope (Leica Microsystems, Wetzlar, Germany) equipped with X100 HCX PL APO NA 1.4 objective or X40 HCX PL APO NA 0.75 or X63 PL APO NA 0.60. Laser excitation line, intensity and emission were chosen according to fluorophores. For AR imaging, lamp and laser were used with the same power intensity. Fibre typing was analysed in 10 μm quadriceps cryosections by immunofluorescence using combinations of the following monoclonal antibodies distributed by DSHB: BA-D5 (MyHC-I; 1:300), BF-F3(MyHC-IIb; 1:300) and SC-71(MyHC-IIa; 1:300). Myofibers not stained were considered MyHC-IIx. Images were captured using a Leica DFC300-FX digital charge-coupled device camera by using Leica DC Viewer software, and morphometric analyses were made using ImageJ.

### Quantitative real-time PCR analysis

Total RNA was extracted with TRIzol (Thermo Fisher Scientific), and RNA was reverse-transcribed using the iScript Reverse Transcription Supermix (1708841 Bio-Rad) following the manufacturer’s instructions. Gene expression was measured by RT-qPCR using the SsoAdvanced Universal Sybr green supermix (1725274 Bio-rad) and the C1000 Touch Thermal Cycler–CFX96 Real-Time System (Bio-Rad). The list of specific primers (Eurofins) is provided in **Supplementary Table 2**. The following TaqMan probes (Applied Biosciences) were used: CACNA1S (Mm00489257), CASQ1 (Mm00486725), CASQ2 (Mm00486742), ATP2A1 (Mm01275320), ATP2A2 (Mm01201431), PV (Mm00443100), SLN (Mm00481536), RYR1 (Mm01175211). Gene expression for castration and ASO experiments was normalized to Cpsf2 (Mm00489754). Gene expression for AR100Q, AR113Q, and doxAR00Q was normalized to Actin (Mm01333821).

### Intracellular Ca^2+^ measurements

Single FDB fibers were isolated and transfected as previously described (78). The transients of fura-2 ratio R were recorded during 2 seconds of stimulations at 60 Hz, with a sampling rate of 1 ms, along with the excitation signals. Data analysis were automated with an ad-hoc Matlab (R) script. After interpolation of the raw data using the built-in function interp1, the following information was extracted. Basal levels were defined as the average of R in a 500 ms time window before the first impulse of the 60-Hz train. Since different cells required slightly different number of stimuli to reach the steady-state of R, we defined the “first peak” as the maximum value within the first five impulses. The amount of Ca^++^ released to cytosolic space was estimated through the integral of R within the first 100 stimuli (about 1.6 seconds). For the transient decay after the last stimulus, R was fitted with a single exponential function, in a time-window of 600-ms, using the built-in Curve Fitting toolbox, and the inverse of the rate of decay (the time constant) was reported in the text. Even if the noise in each trace between two stimuli was too high to give some information, yet the recording of the stimuli in time in our set-up allowed for the aligning of each stimulus at its onset to obtain their ensemble average. The differences in the increments of the ensemble averages from each cell showed a statistical significance between groups, and the different behavior can be observed with the mean value of the ensemble averages for each group.

### *In vivo* ChIP assay

ChIP assay was performed in gastrocnemius muscle using Magna ChIP A/G Chromatin Immunoprecipitation Kit (Millipore) (79). Sheared crosslinked chromatin was immunoprecipitated separately with an anti-AR antibody (3 μg, 17-10489 ChIPAb + androgen receptor Assay Kit, Millipore) or an equal amount of normal IgG (3 μg, androgen receptor Assay Kit, Millipore). After decrosslinking and DNA purification, samples were subjected to quantitative RT–PCR. The oligonucleotide primers used are shown in **Supplementary Table 3**. The regions of amplification contain an androgen-responsive element (ARE)-binding sites for the promoter studied.

### Statistical analysis

To compare measures across groups student’s two sample t-tests and one-way analysis of variance (ANOVA) tests followed by Tukey’s Honest Significant difference post-hoc tests were used for two and more than two group comparisons, respectively. To evaluate body weight and behavioral differences across genotype groups over time two way-ANOVAs with genotype and time as predictor variables, followed by a Tukey’s honest significant difference post-hoc tests was used. Statistical comparison of Kaplan Meier survival curves was performed using the log-rank test. For all tests, significance threshold was set at p < 0.05.

## Acknowledgements

We thank Carlo Reggiani for insightful discussion. We thank Sergio Robbiati, Ludovico Scenna, and Marta Tarter (CIBIO, University of Trento) for help with animals, Alessandro Martino, Elisabetta Broseghini and Piero Rigo (University of Trento) for help with experiments, and Cinzia Ferri (San Raffaele, Milan) for help with peripheral nerve preparation for microscopy. This work was supported by Telethon-Italy and Provincia Autonoma di Trento (TCP12013 to M.Pe.), Telethon-Italy (GGP14147 to D’A.M.), Association Française contre les Myopathies (18722 to G.S. and M.Pe., and 22221 to M.P and M.B.), Muscular Dystrophy Association (479363 to M.Pe.), Kennedy’s Disease Association Research Grant (U-GOV PIRA_EPPR19_01 to M.Pi) Marie Curie International Outgoing Fellowships (PIOF-GA-2011-300723 to S.P.), Akira Arimura Foundation (E.Z.), University of Padova (Bird project), Italian Ministry of Health (RF-2011-02350097 to G.S. and M.Pe.), U. S. National Institutes of Health (R01 NS055746 to A.P.L.), and Fondazione Umberto Veronesi Fellowship (to M.C. & E.Z.). S.N. is supported by a Rackham Pre-doctoral Fellowship and C.R. is supported by a Wellcome Trust Clinical Research Career Development Fellowship (205162/Z/16/Z). The Neuromuscular Bank of Tissues and DNA samples, member of the Telethon Network of Genetic Biobanks (project no. GTB12001), funded by Telethon Italy, and of the EuroBioBank network, provided us with biopsy specimens.

## Author contribution

MC, CM, MPi, and MPe contributed to study design and performed GCN, behavior studies, histological analysis, biochemistry experiments, gene expression analysis. CS, DP, MJP, EZ contributed to transgenic mouse line generation and phenotype assessment. NS and LAP performed experiments on knock-in mice. NL, MC, ML, and BB performed analysis of muscle force and Ca^++^ dynamics. AA performed ChIP experiment. RV performed TMRM assay. SP genotyped animals from July 2013 through spring 2014. D’AM contributed to peripheral nerve pathological analysis. FS performed statistical analysis. EP, GS, CR, MS contributed to data interpretation. ED performed the bioinformatical analysis. MB performed analysis of gene expression, contributed to study design, and data interpretation. EZ, CM, MPi, and MPe wrote the manuscript.

