## Supplementary material for "Polyglutamine-expanded androgen receptor disrupts muscle triad, calcium dynamics and the excitation-contraction coupling gene expression program"

**Supplementary Table 1**

**ARE sequences in the regulatory elements of the indicated genes**

| gene name | start | end | sequence | P value |
|---|---|---|---|---|
| Cacna1s | -4903 | -4887 | AGGAACTCCATGCCCCA | 4.40E-04 |
| Cacna1s | -4320 | -4304 | TGAGAAATTGTGTTCCT | 6.09E-04 |
| Cacna1s | -3637 | -3621 | ACGAACACTGTGAACAC | 1.93E-04 |
| Cacna1s | -3359 | -3343 | AGGTACAGAGTGAGTTC | 2.36E-04 |
| Cacna1s | -3202 | -3186 | GAGACCCATATGTCCTC | 6.84E-04 |
| Cacna1s | -2956 | -2940 | GTGGACACAGTGCTCTT | 1.92E-04 |
| Cacna1s | -2162 | -2146 | AGATACCTGCTGTGCCT | 2.85E-04 |
| Cacna1s | -1567 | -1551 | CGCTACATCAAGTACCT | 9.26E-04 |
| Cacna1s | 363 | 379 | AGGTACGGGGTGGTCAC | 5.78E-04 |
| Cacna1s | 419 | 435 | GGGAAAGGGGTGAACCT | 1.92E-04 |
| Atp2a1 | -4496 | -4480 | AGGAAGAGCTGGTCCCC | 8.63E-04 |
| Atp2a1 | -3695 | -3679 | GGTTCCAGCATGTTCTG | 5.29E-04 |
| Atp2a1 | -1704 | -1688 | AGGAACACCTTGTCCTG | 4.14E-06 |
| Atp2a1 | -437 | -421 | GAGAACTTCCTCTGCTA | 1.05E-03 |
| Ryr1 | -3207 | -3191 | GGGCAAGCAGTCTACTA | 9.12E-04 |
| Ryr1 | -3090 | -3074 | GAGGGCATCAGGTTCCT | 5.28E-04 |
| Ryr1 | -3003 | -2987 | GAGCAGCAGGTGTTCTT | 2.97E-04 |
| Ryr1 | -1559 | -1543 | GGGGAAATTGTGCCCCC | 8.05E-04 |
| Ryr1 | -483 | -467 | GGGGATGTGGTGTGCAT | 5.14E-04 |
| Atp2a2 | -4320 | -4304 | AAGAACACTGGCTGCTT | 4.78E-04 |
| Atp2a2 | -3223 | -3207 | GAAAACAGACTGAGCCC | 5.70E-04 |
| Atp2a2 | -2487 | -2471 | AGGAGCAGTCAGTGCTC | 5.89E-04 |
| Atp2a2 | -643 | -627 | TGGGATGCGGTGTTCCC | 7.89E-05 |
| Casq1 | -3949 | -3933 | GAGAAGGAGACGTACTA | 2.57E-04 |
| Casq1 | -3737 | -3721 | AAGCCCAGCCTGTTCTG | 1.67E-04 |
| Casq1 | -3643 | -3627 | GGTTACTTGGTGTGCAG | 1.07E-03 |
| Casq1 | -2997 | -2981 | GGGCAGACTGTGATTCC | 9.37E-04 |
| Casq1 | -2882 | -2866 | AGGCAAATAATGTATCA | 4.13E-04 |
| Casq1 | -1047 | -1031 | GGGCACAGGGTCCACCT | 3.70E-04 |
| Casq1 | 52 | 68 | TGGCCCACTCTCTACCC | 8.88E-04 |
| Casq1 | 157 | 173 | AGGCCCAGATTGTACTA | 1.13E-04 |
| Casq1 | 332 | 348 | GTGAACGCCAAGAACTA | 9.77E-04 |
| Pln | -4106 | -4090 | AATTACAAAATGTGCCT | 8.89E-05 |
| Pln | -3650 | -3634 | GAGAACACAATGAAGCA | 5.70E-04 |
| Pln | -2829 | -2813 | TGGCACATTATTTTCAC | 5.30E-04 |
| Pln | -2516 | -2500 | ATGAACAGTGCTTTCCT | 6.40E-04 |
| Pln | -2300 | -2284 | AAGGACACTGAGCACTA | 3.64E-04 |
| Pln | -1147 | -1131 | AGGCACACAGTTTTCAT | 1.85E-04 |
| Pln | -123 | -107 | AAGCACAAAGTGTTAAT | 2.17E-04 |
| Pln | -58 | -42 | AAGAACATGGCTAACCA | 6.80E-04 |

| | | | | |
|---|---|---|---|---|
| Casq2 | -4832 | -4816 | GGTAACAGATGGTTCCC | 3.79E-04 |
| Casq2 | -4421 | -4405 | CAGGACAGGGGGTTCAA | 4.95E-04 |
| Casq2 | -2635 | -2619 | GGGTACCCACAGACCCT | 1.05E-03 |
| Casq2 | -1118 | -1102 | TGGTACAGGGTCCGCCT | 8.19E-04 |
| Sln | -4281 | -4265 | GATCACACTATGTAATC | 4.21E-04 |
| Sln | -4119 | -4103 | AGGGACATAGGTTACTG | 1.02E-03 |
| Sln | -3612 | -3596 | AGTCACAGAGTGACCTC | 7.21E-04 |
| Sln | -3125 | -3109 | GGGAAAGAAGTGTTATA | 7.44E-04 |
| Sln | -1427 | -1411 | GGTTACAGGGTTTGCCA | 8.97E-04 |
| Sln | -860 | -844 | GAGCACTAGGTGTTCCA | 2.57E-05 |

**Supplementary table 2**

**Primers Real Time used for quantitative PCR analyses**

| mouse | Forward (5'-3') | Reverse (5'-3') |
|---|---|---|
| CACNA1S | CAGACACAGAGAGCTTGTATGAA | TTCACTAGGTCATGGCACTTC |
| CASQ1 | CCC TGT AGA GTT GAT TGA AGG TGA ACG | CCT CGT AGG CTT TGT AAT GCT CTG AGT C |
| CASQ2 | CCAGCTGAAGGAGATTGTACTG | AGCAAGCTTGGCCTCTTT |
| ATP2A1 | CCACCAACCAGATGTCAGTT | TCCCTCAGGAGCATAAGTAGAG |
| ATP2A2 | CTTATCTTGGTAGCCAATGCAATC | TTGCCCATTTCAGGCTCATA |
| PV | AGGTGAAGAAGGTGTTCCATATT | TCTCTGGCATCTGAGGAGGAA |
| SLN | GCTCCTCTTCAGGAAGTGAAG | TGGCCCCTCAGTATTGGTAGG |
| RYR1 | GCAACCGCCTTTGCTTTC | ACTGACAGAGACTGCTCCA |
| B-ACTIN | GACAGGATGCAGAAGGAGATTACTG | CTCAGGAGGAGCAATGATCTTGAT |
| FOXO3A | CGCTGTGTCCCTACTTCA | CCCGTGCCTTCATTCTGA |
| LC3B | CACTGCTCTGTCTTGTGTAGGTTG | TCGTTGTGCCTTTATTAGTGCATC |
| MUSA1 | TCGTGGAATGGTAATCTTGC | CCTCCCGTTTCTCTATCACG |
| SMART | TCAATAACCTCAAGGCGTTC | GTTTTGCACACAAGCTCCA |
| SQSTM | CCCAGTGTCTTGGCATTCTT | AGGGAAAGCAGAGGAAGCTC |
| MYOG | CTTGCTCAGCTCCCTCAAC | TGGGAGTTGCATTCACTGG |
| MYH3 | GGGACCTTGCCAAGAAGAA | GTCGTTCCTCACGGTCTTG |
| MUSK | ATCACCACGCCTCTTGAAAC | TGTCTTCCACGCTCAGAATC |
| MYH8 | GAGGGCATCCGCATCTG | GATGAACTGTCCCTCTGGAATAG |
| KEAP1 | CATCCACCCTAAGGTCATGGA | GACAGGTTGAAGAACTCCTCC |
| NFE2L1 | CAGCAGTGGCAAGATCTCAT | GGGCATTGTACAGAATCTCACT |
| NFE2L2 | CAGGAGAATTCCTCCCAATTC | GGCATCTTGTTTGGGAATGTG |
| SDHA | TTACCTGCGTTTCCCCTCAT | AAGTCTGGCGCAACTCAATC |
| NQO1 | AGTGCTCGTAGCAGGATTTG | AGTGGTGATAGAAAGCAAGGT |
| PPARGC1A | GGAATGCACCGTAAATCTGC | TTCTCAAGAGCAGCGAAAGC |

**Supplementary table 3**

**Primers used for ChIP analysis**

| mouse | Forward (5'-3') | Reverse (5'-3') |
|---|---|---|
| SLN | ACTCGTCCTTGTCTGATTTGAG | CACTGTGTGGGTACTAGGATT |

### Supplementary Figure legends

##### Supplementary Figure 1

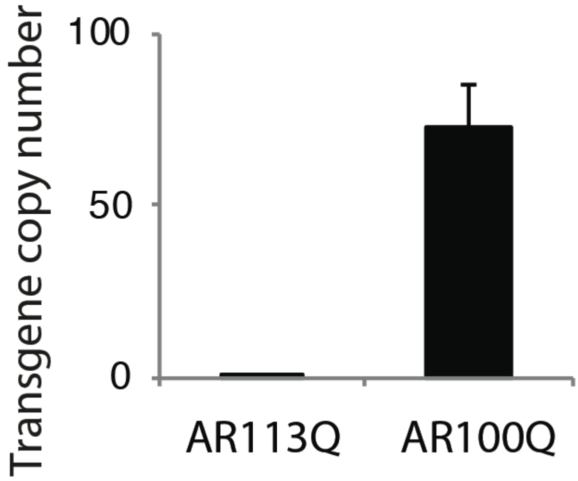


**Supplementary Figure 1. Copy number of *hAR* transgene in AR100Q mice.**
Analysis of gene copy number by quantitative PCR. AR113Q knock-in mice, in which the endogenous *AR* exon 1 was replaced with the human exon 1, were used as reference and normalized to 1. Graph, mean ± sem, n = 3.
Gene copy number (GCN) was evaluated using RT-qPCR and the qbase+ software (1). Genomic DNA was extracted from 5 mice of each line using ReliaPrep™ gDNA Tissue Miniprep System (Promega). Specific primers were designed to amplify the hAR transgene and two housekeeping genes with a known GCN=2: GUSB (Gusb glucuronidase, beta *mus musculus*), and TfR (Transferrin receptor *mus musculus*). A mouse sample was used as an inter-run control and the transgenic mice were compared to the male knock-in mouse gDNA where GCN=1 for hsaAR (2). Primers were designed as followed: hsaAR Forward 5'- CTTCACCGCACCTGATGTG, hsaAR Reverse 5,- TAAGGTCCGGAGTAGCTATC, mmuGusb Forward 5'- CCTGGATGTCCGGGTGAATC, mmuGusb Reverse 5'- GAGGTATGTGCACCGGGATG, mmuTfR Forward 5'- ATACCCCAAATTTTGACCAGCC, mmuTfR Reverse 5'- GACCTTGCCTCAAAGAAAAACCT.

**Supplementary Figure 2**

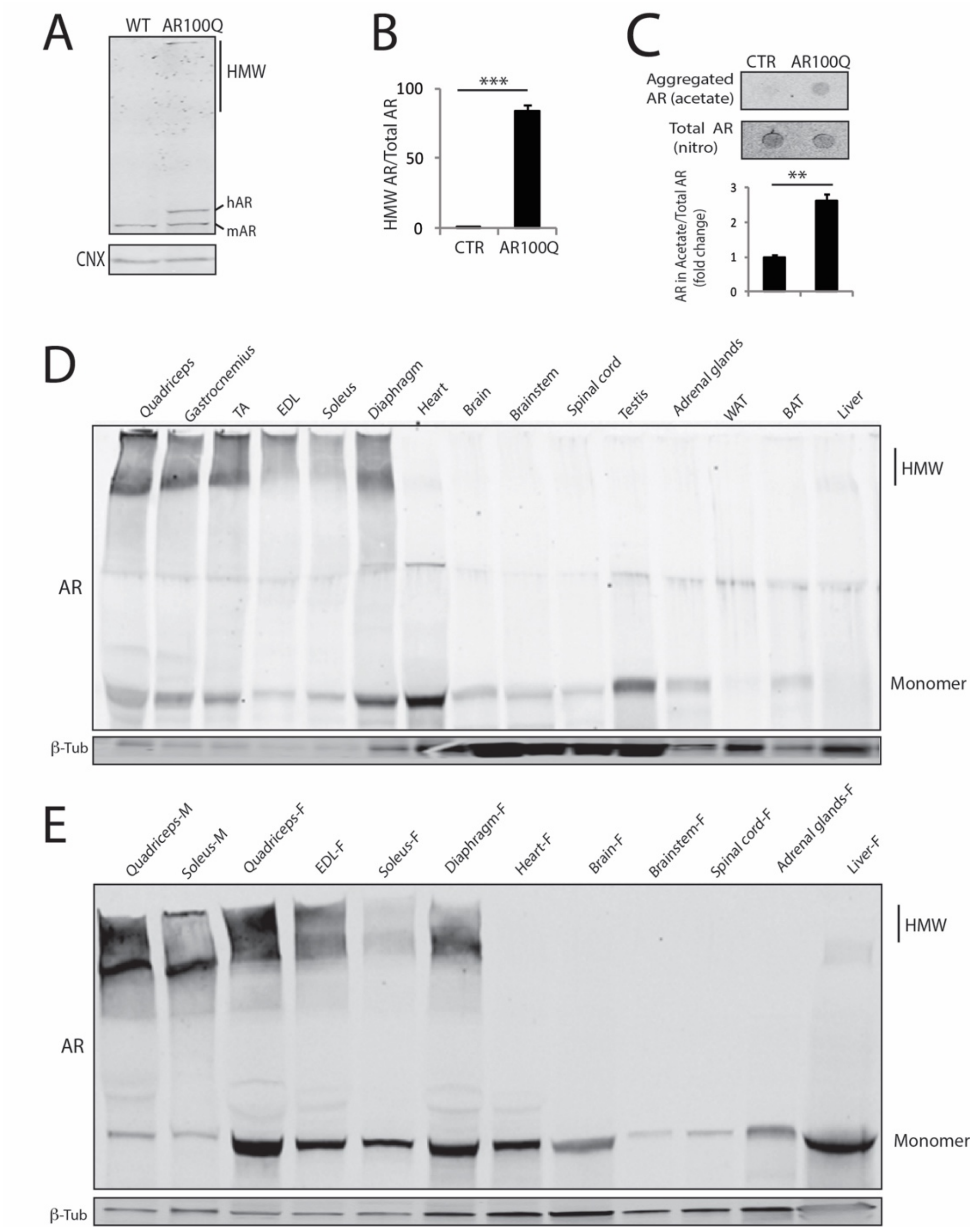


**Supplementary Figure 2. PolyQ-expanded AR forms aggregates selectively in skeletal muscle.**

A) Western blotting analysis of AR aggregation in the brainstem of 8-week-old control (WT) and AR100Q mice. Proteins lysates were performed in RIPA buffer containing high SDS (2%).

B) Quantification of AR high-molecular weight (HMW) species in the skeletal muscle of 8-week-old control (CTR) and AR100Q mice (n = 3) shown in Figure 1A (main text).

C) Filter retardation assay of protein extracts from quadriceps muscle of 8-week-old control (CTR) and AR100Q mice (n = 3). Quantification is shown at the bottom panel.

D) Western blotting analysis of AR aggregation in the indicated tissues of 8-week-old AR100Q mice (n = 3).

E) Western blotting analysis of AR aggregation in 8-week-old male (M) and female (F) AR100Q mice (n = 3).

AR was detected with a specific antibody, and calnexin (CNX) and beta-tubulin (β-Tub) were used as loading controls. mAR, mouse endogenous AR; hAR, human AR transgene. Shown is one experiment representative of at least three experiments performed on different mice/genotype. Graph, mean ± sem, n = 3 mice per group, student's t test, ** p = 0.01, *** p = 0.001.

**Supplementary Figure 3**

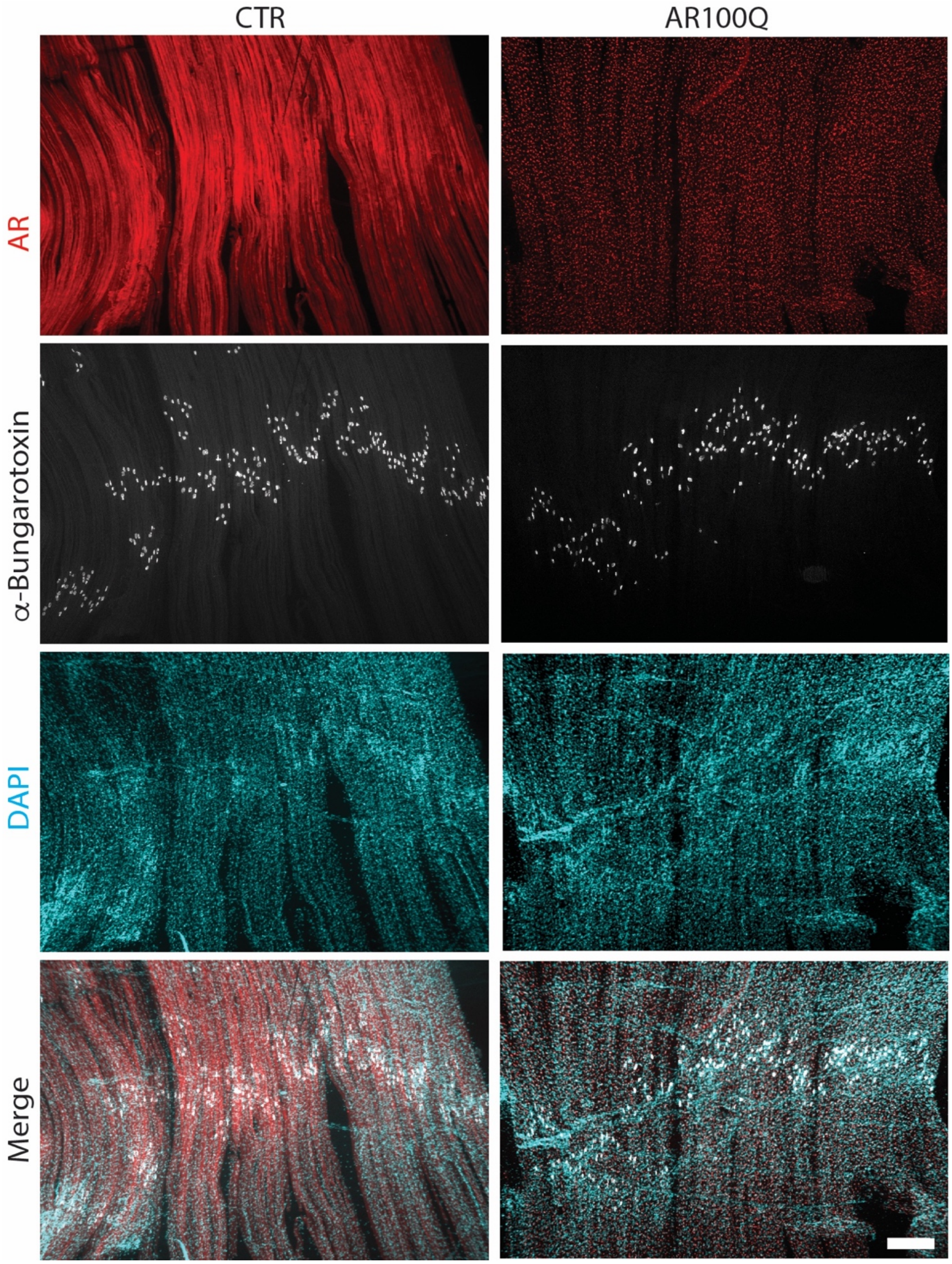


**Supplementary Figure 3. Expression of polyQ-expanded AR resulted in age-dependent inclusion body formation in the muscle of AR100Q mice.**
Immunofluorescence analysis of AR subcellular localization in intact fibers from levator auris longus (LAL) muscle of 8-week-old control (CTR) and AR100Q mice. AR was detected with a specific antibody, NMJs by staining with α-Bungarotoxin, and nuclei with DAPI. Shown are representative images of at least 3 mice for each group. Bar, 250 micron.

**Supplementary Figure 4**

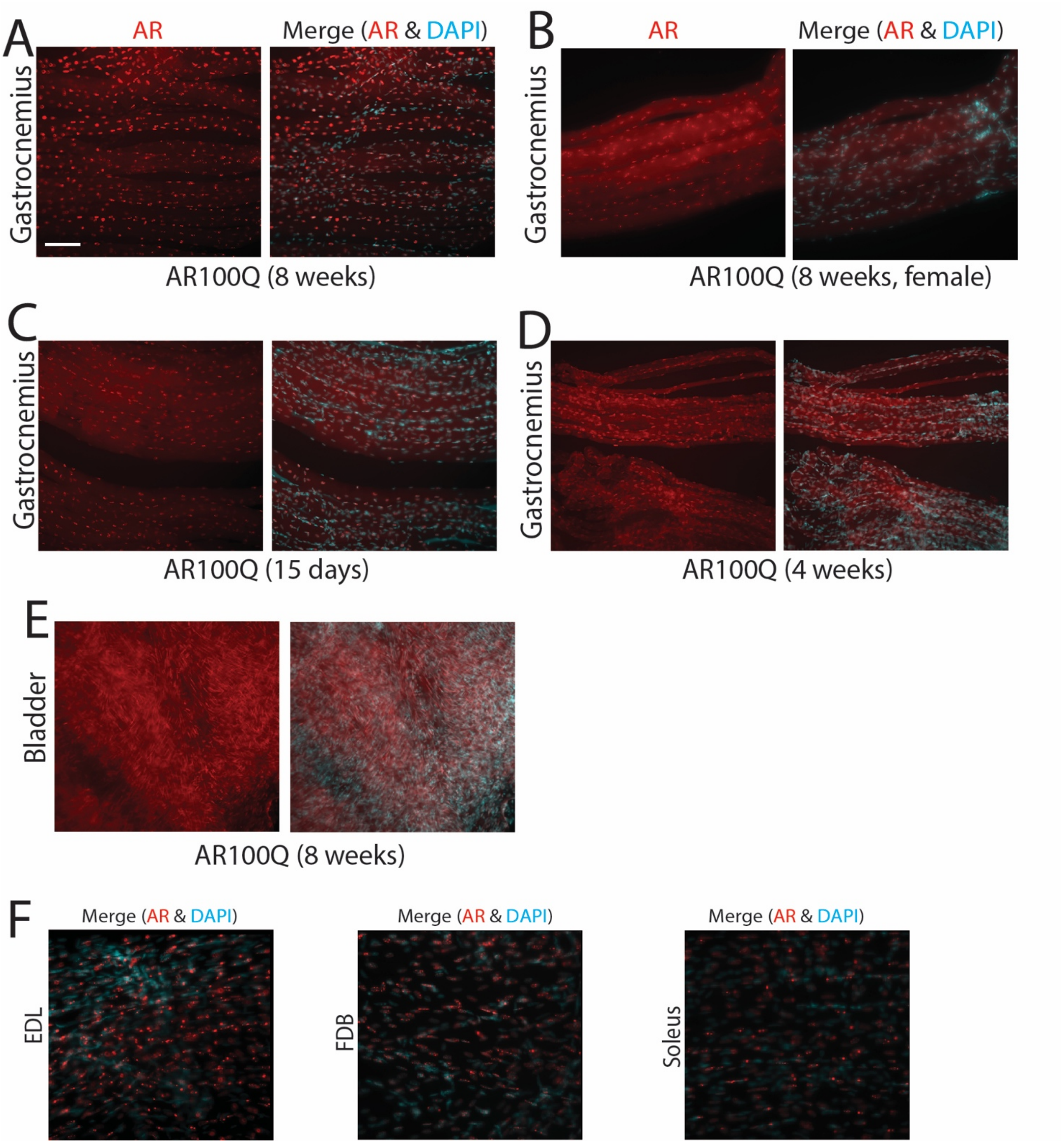


**Supplementary Figure 4. Inclusion body formation in the muscle of AR100Q was higher in male compared to female mice and in fast muscles compared to slow muscles.**

A-F) Immunofluorescence analysis of AR subcellular localization in intact fibers from the indicated muscles of AR100Q mice. Bar, 100 micron.

AR was detected with a specific antibody and nuclei with DAPI. Shown are representative images of at least 3 mice for each group.

**Supplementary Figure 5**

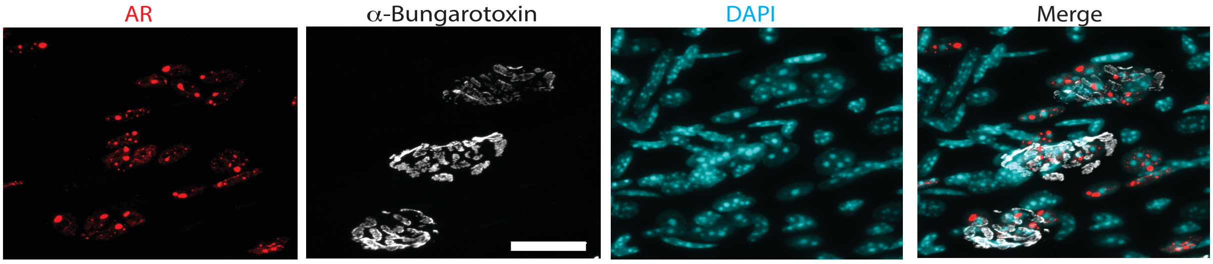


**Supplementary Figure 5. Inclusion body formation in the subsynaptic nuclei of muscle of AR100Q mice.** Immunofluorescence analysis of AR subcellular localization in intact fibers from LAL muscle of 8-week-old AR100Q mice. AR was detected with a specific antibody, NMJs by staining with alpha-Bungarotoxin, and nuclei with DAPI. Shown are representative images of at least 3 mice. Bar, 25 micron.

**Supplementary Figure 6**

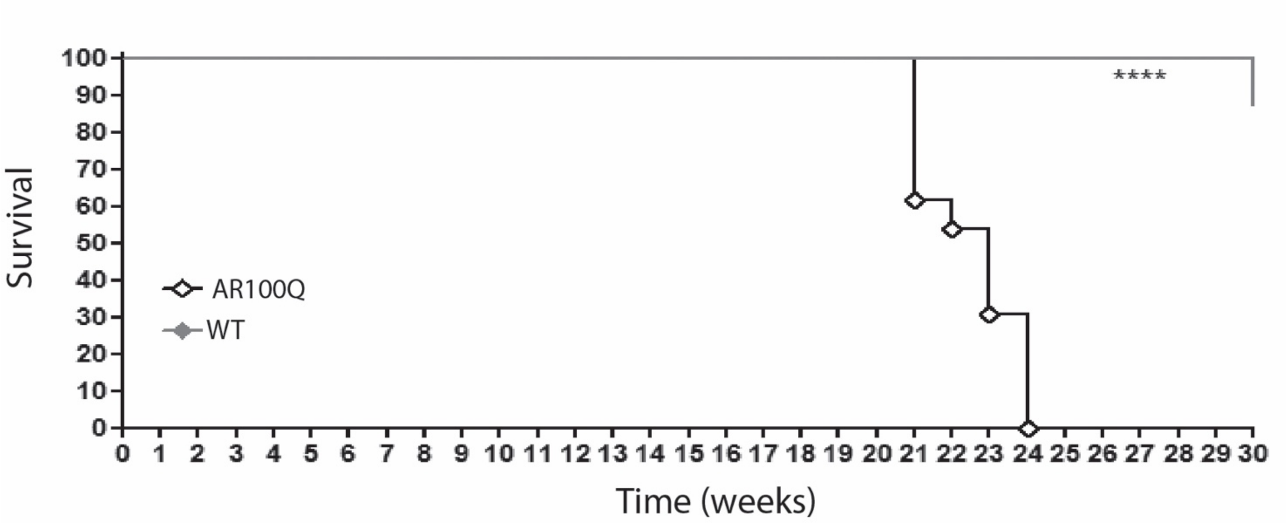


**Supplementary Figure 6. Decreased lifespan of female AR100Q mice.**
Kaplan-Meier analysis of lifespan of WT (n = 13) and AR100Q (n = 13) female mice. Survival analysis revealed that female AR100Q mice have a median survival of 23 weeks. Survival curves were compared using Log-rank (Mantel-Cox) test.

**Supplementary Figure 7**

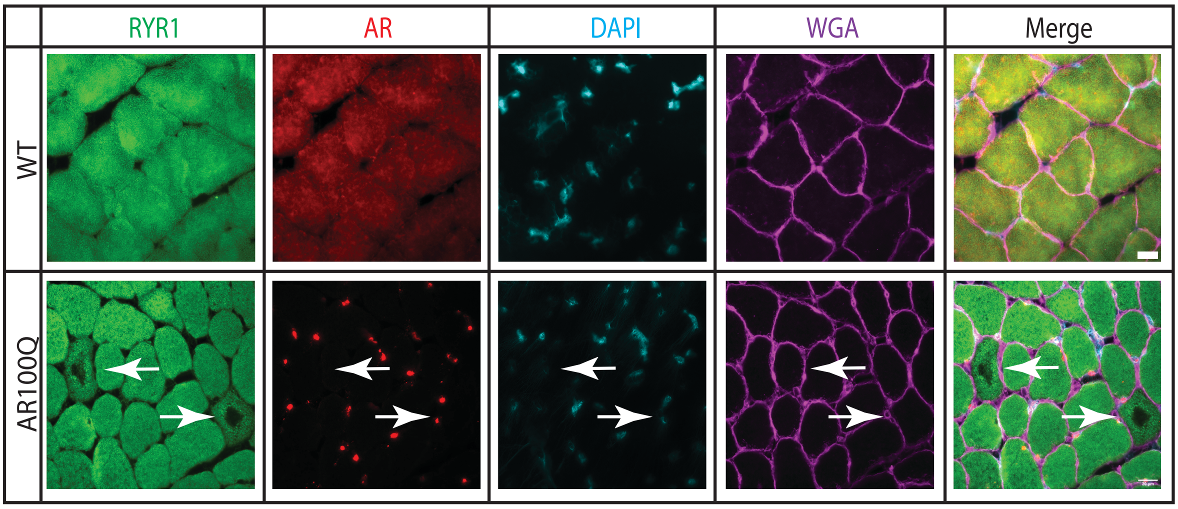


**Supplementary Figure 7. Mitochondrial pathology in the muscle of AR100Q mice.**

Immunofluorescence analysis of RYR1 (green), AR (red), nuclei (DAPI, blue), and wheat germ agglutinin (WGA) lectin (purple) in the quadriceps of 8-week-old WT and AR100Q mice (n = 3). Shown are representative images. Bar, 20 micron.

**Supplementary Figure 8**

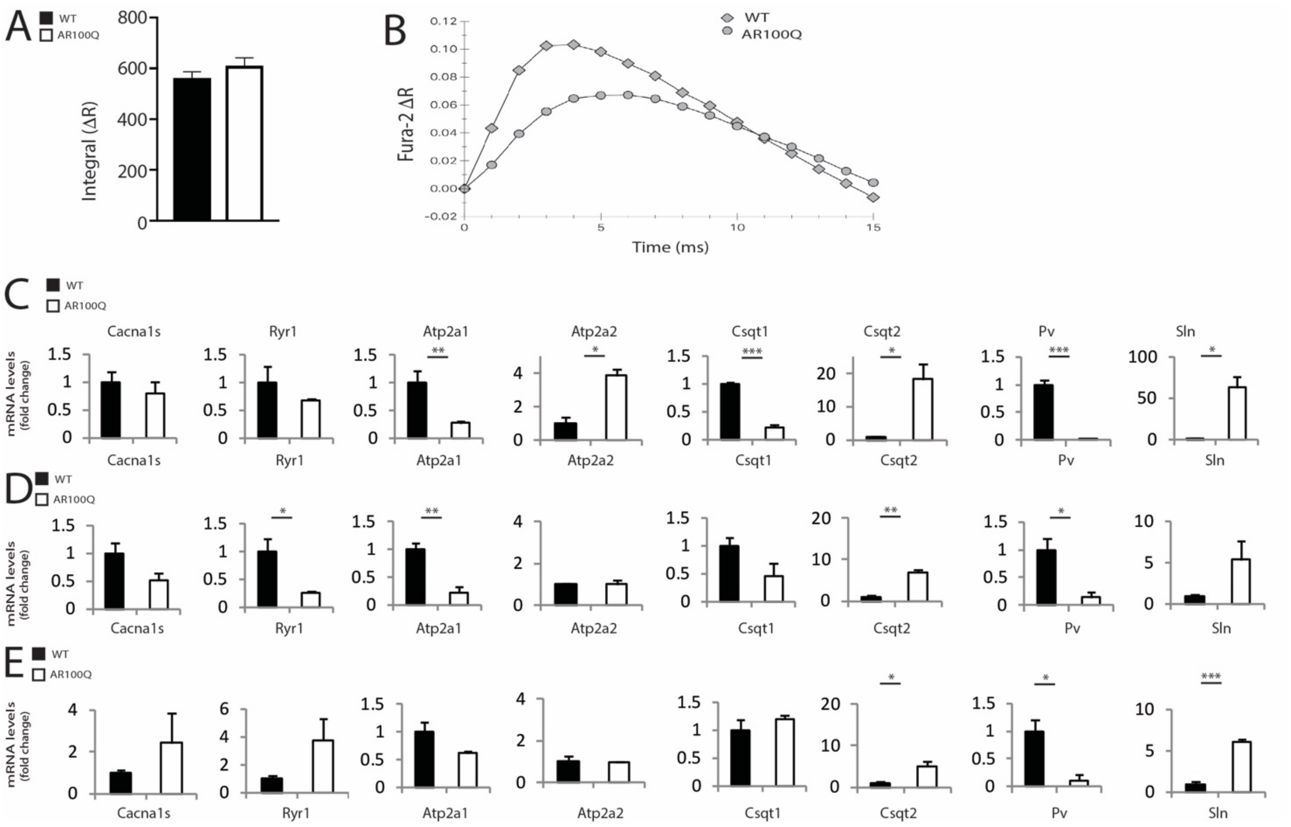


**Supplementary Figure 8. Expression of ECC genes is disrupted in fast-twitch muscles of AR100Q mice.**

A-B) Analysis of $Ca^{++}$ levels in response to tetanic stimulation in fibers isolated from FDB of 8-week-old WT and AR100Q mice (n = 3). The amount of calcium accumulated in the cytosol during the tetanic stimulation (A), calculated as the integral of the FURA_2 ratio in one second, did not show significant differences between WT and AR100Q mice. The representative ensemble average of the $Ca^{++}$ transients between consecutive stimuli (B) showed a faster increase as well as a faster decrease in WT compared to AR100Q mice.

C-E) Real-time PCR analysis of the indicated genes in the EDL (C), FDB (D), and soleus (E) muscles of 8-week-old WT and AR100Q mice. Graph, mean ± sem, n = 3-5 mice for each genotype, student's t test, * p = 0.05, ** p = 0.01, *** p = 0.001.

**Supplementary Figure 9**

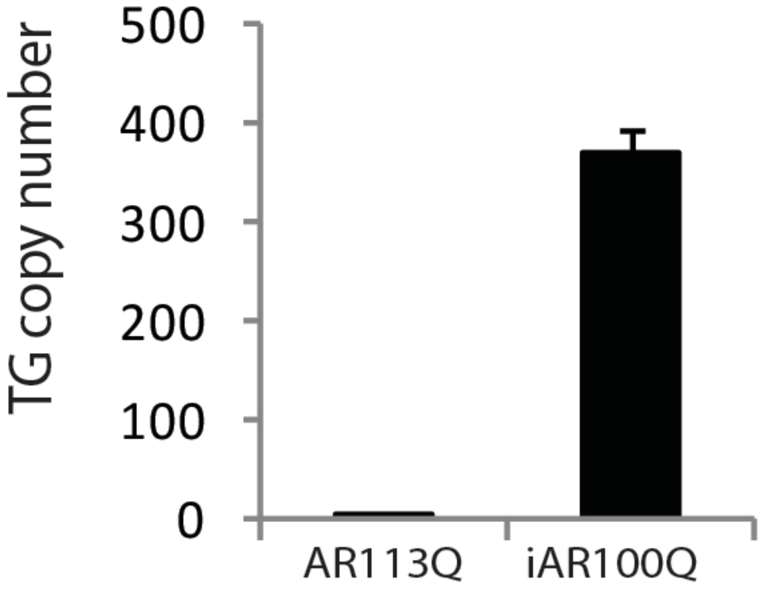


**Supplementary Figure 9. Copy number of *hAR* transgene in iAR100Q mice.**
Analysis of gene copy number by quantitative PCR. AR113Q knock-in mice, in which the endogenous *AR* exon 1 was replaced with the human exon 1, were used as reference and normalized to 1. Graph, mean ± sem, n = 3, student's t test. Methods are described in Supplementary Figure 1 caption.

**Supplementary Figure 10**

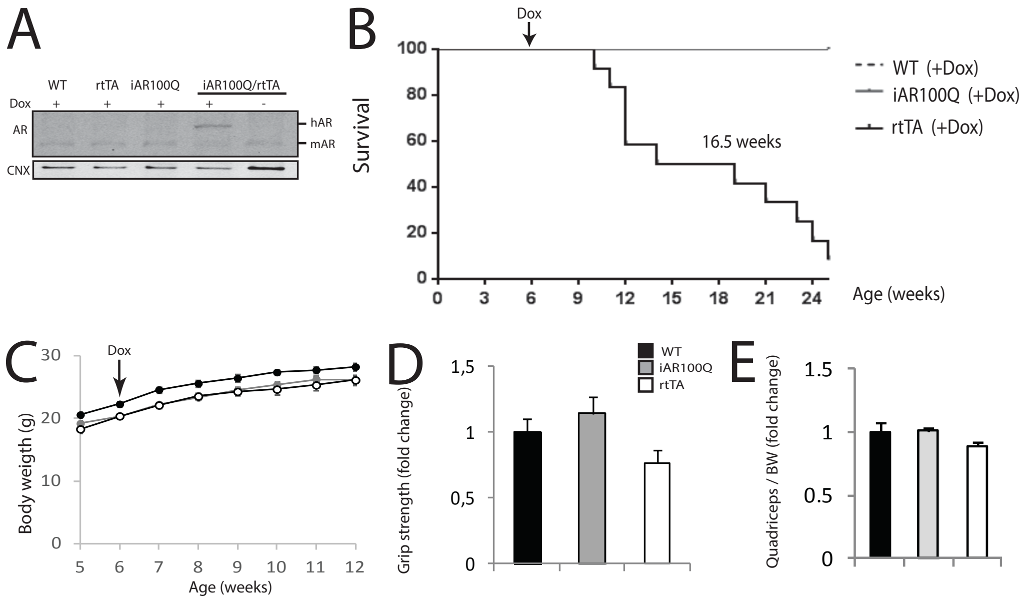


**Supplementary Figure 10. Generation and characterization of inducible transgenic mice expressing AR100Q.**

A) Western blotting analysis of AR levels in WT (n= 3), rtTA (n= 3), iAR100Q (n= 3), iAR100Q/rtTA (n=3) mice treated with either vehicle (sucrose 50g/l) or doxycycline (sucrose 50g/l, dox=1g/l).

B) Kaplan-Meier survival curves of WT (n = 8), rtTA (n =12), and iAR100Q (n = 8) mice treated with either vehicle or dox. Survival curves were compared using Log-rank (Mantel-Cox) test. Although dox treatment reduced the median survival of the rtTA mice to 16.5 weeks, the effect of induction of expression of AR100Q on survival was statistically different from that of rtTA.

C) Temporal changes in mean BW of WT (n = 8), rtTA (n = 8), and iAR100Q (n = 8) mice treated with either vehicle or dox.

D) Grip strength analysis of muscle force of WT (n = 8), rtTA (n =8), and iAR100Q (n = 8) mice treated with either vehicle or dox.

E) Quadriceps weight normalized to BW of WT (n = 5), rtTA (n = 5), and iAR100Q (n =5) mice treated with either vehicle or dox.

Graph, mean ± sem, (D and E) one-way ANOVA followed by Tukey's Honest Significant difference post hoc test.
